# Chromosome-scale assemblies of Acanthamoeba castellanii genomes provide insights into Legionella pneumophila infection-related chromatin re-organization

**DOI:** 10.1101/2021.10.26.465878

**Authors:** Cyril Matthey-Doret, Morgan J. Colp, Pedro Escoll, Agnès Thierry, Bruce Curtis, Matt Sarrasin, Michael W. Gray, B. Franz Lang, John M. Archibald, Carmen Buchrieser, Romain Koszul

## Abstract

The unicellular amoeba *Acanthamoeba castellanii* is ubiquitous in aquatic environments, where it preys on bacteria. The organism also hosts bacterial endosymbionts, some of which are parasitic, including human pathogens such as *Chlamydia* and *Legionella* spp. Here we report complete, high quality genome sequences for two extensively studied *A. castellanii* strains, Neff and C3. Combining long- and short-read data with Hi-C, we generated near chromosome-level assemblies for both strains with 90% of the genome contained in 29 scaffolds for the Neff strain and 31 for the C3 strain. Comparative genomics revealed strain-specific functional enrichment, most notably genes related to signal transduction in the C3 strain, and to viral replication in Neff. Furthermore, we characterized the spatial organization of the *A. castellanii* genome and showed that it is reorganized during infection by *Legionella pneumophila.* Infection-dependent chromatin loops were found to be enriched in genes for signal transduction and phosphorylation processes. In genomic regions where chromatin organization changed during *Legionella* infection, we found functional enrichment for genes associated with metabolism, organelle assembly, and cytoskeleton organization, suggesting that changes in chromosomal folding are associated with host cell biology during infection.

## Introduction

The first amoebae were isolated in 1913 [1], and the genus *Acanthamoeba* was established in 1931 by Volkonsky [2]. It comprises different species of free living, aerobic, unicellular protozoa, present throughout the world in soil and nearly all aquatic environments [3]. The life cycle of *Acanthamoeba* includes a dormant cyst with minimal metabolic activities under harsh conditions and a motile trophozoite that can feed on small organisms and reproduce by binary fission in optimal conditions [4]. *Acanthamoeba* is perhaps most widely known from its role as a human pathogen, acting to cause the vision-threatening eye infection *Acanthamoeba* keratitis, but it can also cause serious infections of the lungs, sinuses, skin, and a central nervous system disease called granulomatous amoebic encephalitis [5]. The species *Acanthamoeba castellanii* was first isolated in 1930 by Castellani as a contaminant of a yeast culture [6].

In their natural environment, *Acanthamoeba* spp. are characterized by the ability to change their shape through pseudopode formation and are considered professional phagocytes as they feed on bacteria, but may also phagocytose yeasts and algae. However, some bacteria are resistant to degradation and live as endosymbionts in these protozoa, and others even use the amoeba as a replication niche. Thus *Acanthamoeba* are also reservoirs of microorganisms and viruses, including human pathogens, which have adapted to survive inside these cells and resist digestion, persist or even replicate as intracellular parasites. At least 15 different bacterial species, two archaea and several eukaryotes and viruses have been shown to interact with *Acanthamoeba* in the environment and may even co-exist at the same time within the same host cell [7].

Although it was observed early on that bacteria could resist digestion of free-living amoebae [8], it was not until the discovery that *Legionella pneumophila* replicated in amoebae that researchers began studying the bacterium-amoeba relationship in depth [9]. *L. pneumophila* is the agent responsible for Legionnaires’ disease, a severe pneumonia that can be fatal if not treated promptly. In addition, many species of amoebae have the ability to form highly resistant cysts in hostile environments, providing shelter for their intracellular parasites [10]. Indeed, it is thought that *L. pneumophila* may survive water disinfection treatments and contaminate water distribution systems by encystation [11, 12, 13]. From these contaminated water sources, *L. pneumophila* can reach the human lungs via aerosols contaminated with the bacteria and replicate within the alveolar macrophages that are, like amoebae, phagocytic cells.

*L. pneumophila* has the ability to escape the lysosomal degradation pathway of both *A. castellanii* and human alveolar macrophages through the formation of a protective vacuole (the Legionella-containing vacuole or LCV) where it multiplies to high numbers. Once the host cell has been fully exploited and nutrients become limited, *L. pneumophila* exits the host and infects a new cell [14].

To establish the LCV and replicate, *L. pneumophila* secretes over 300 effector proteins into the host cytoplasm via a type four secretion system (T4SS) called Dot/Icm [15], thereby manipulating host pathways and redirecting nutrients to the LCV [16, 17]. In the early stages of infection, many of these proteins target the host secretory pathway, including several small GTPases, to recruit endoplasmic reticulum-derived vesicles to the LCV [18]. During the intracellular cycle, a wider range of processes, including membrane trafficking, cytoskeleton dynamics, and signal transduction pathways, are targeted by these effectors [19, 20]. *L. pneumophila* also directly alters the genome of its host by modifying epigenetic marks of the host genome in human macrophages and *A. castellanii*. It secretes an effector named RomA with histone methyltransferase activity that is targeted to the nucleus. RomA carries out genome-wide trimethylation of K14 of histone H3 [21], leading to transcriptional changes that modulate the host response in favor of bacterial survival [21]. Concomitantly, *L. pneumophila* infection leads to genome-wide changes in gene expression [22]. In many eukaryotes, gene regulation is intertwined within the three-dimensional organization of chromosomes. The functional interplay between gene regulation and higher-order chromatin elements such as loops, self-interacting domains and active/inactive compartments is actively being studied [23, 24]. Therefore, the infection of *A. castellanii* by *L. pneumophila* provides an amenable model with which to investigate how an intracellular bacterial infection may affect the regulation of chromosome folding, and its consequences, in a eukaryotic host.

The investigation of genome organization and regulation of *A. castellanii* in response to infection requires a highly contiguous genome assembly. The reference genome sequence for *A. castellanii,* NEFF-v1 [25], is based on the Neff strain, isolated from soil in California in 1957 [26]. This assembly is widely used by different laboratories studying *A. castellanii,* but is fragmented into 384 scaffolds comprising 3192 contigs, which makes chromosome-level analyses difficult, if not impossible, and basic features of the *A. castellanii* genome, such as the number of chromosomes and ploidy, remain undetermined. In addition, many teams investigating bacteria-amoeba interactions use the “C3” strain (ATCC 50739), isolated from a drinking water reservoir in Europe in 1994 and identified as a mouse pathogen [27]. However, genomic information is scarce for this strain and little is known about its similarity to the Neff strain. Notably, these two *A. castellanii* strains have been cultivated for several decades and were isolated from different ecological niches, but the extent of conservation between their genomes is unknown. It is difficult to investigate the factors that determine the susceptibility of different *A. castellanii* strains to the pathogen without proper genomic resources. These resources would also be required to apply genome-wide omics approaches.

The goal of this work was to study how the *A. castellanii* C3 strain responds to *L. pneumophila* infection through the lens of the three-dimensional organisation of its genome. This analysis required the generation of a high quality reference genome sequence of the C3 strain, as well as a new and improved assembly of the Neff reference genome. Illumina, Nanopore long read, and Hi-C data were used to generate near chromosome-level assemblies of the genomes of both strains. Surprisingly, the new Neff and C3 assemblies have a (gap-excluded) sequence divergence of 6.7%. We find evidence for strain-specific enrichment of a handful of functions, including ones related to signal transduction in C3, and one relating to viral replication and virion assembly in Neff. Using the C3 assembly, RNA-seq and Hi-C, we were able to analyze the genome folding and expression changes of *A. castellanii* in response to the infection by *L. pneumophila*. We found infection-dependent chromatin loops to be enriched in genes involved in signal transduction and phosphorylation.

## Results

The *A. castellanii* Neff and C3 genome assemblies are highly contiguous and complete We used a combination of Illumina short reads, Oxford Nanopore long reads and Hi-C to assemble each genome to chromosome scale, with 90% of the Neff genome contained within 28 scaffolds. This is in contrast to a previous estimate of approximately 20 chromosomes inferred using pulsed-field gel electrophoresis [28]. For both the Neff and C3 strains, we first generated a raw *de-novo* assembly using Oxford Nanopore long reads. To account for the error prone nature of long reads, we polished the first draft assemblies with paired-end shotgun Illumina sequences using HyPo [29]. The polished assemblies were then scaffolded with long range Hi-C contacts using our probabilistic program instaGRAAL, which exploits a Markov Chain Monte Carlo algorithm to swap DNA segments until the most likely scaffolds are achieved [30]. Following the post-scaffolding polishing step of the program (see [30]), the final genome assemblies displayed better contiguity (Table 1), completion, and mapping statistics than the previous versions, with the cumulative scaffold lengths quickly reaching a plateau (Fig. 1a). The assemblies of both strains are also slightly longer, with a smaller number of contigs than the original Neff assembly (NEFF-v1) (Fig. 1b). The BUSCO-completeness scores for both assemblies are also improved, with 90.6% (Neff) and 91.8% (C3) complete eukaryotic universal single copy orthologs, compared to 77.6% for NEFF-v1. We also noted an increased proportion of properly paired shotgun reads from 71% for NEFF-v1 to 84% for our new Neff assembly, suggesting a reduced number of short mis-assemblies. Hi-C contact maps present a convenient readout to explore large mis-assemblies in genome sequences [31]. While this allowed us to manually address major unambiguous mis-assemblies, a number of visible mis-assemblies remain in complex regions such as repeated sequences near telomeres and ribosomal DNAs (rDNAs). These mis-assemblies could not be resolved with the data generated herein. In the C3 assembly, there are also a few (at least 5) interchromosomal mis-assemblies which appear to be heterozygous and cannot be resolved without a phased genome. We also found shotgun coverage to be highly heterogeneous between scaffolds, which is suggestive of aneuploidy (Fig. S1).

**Figure 1.**
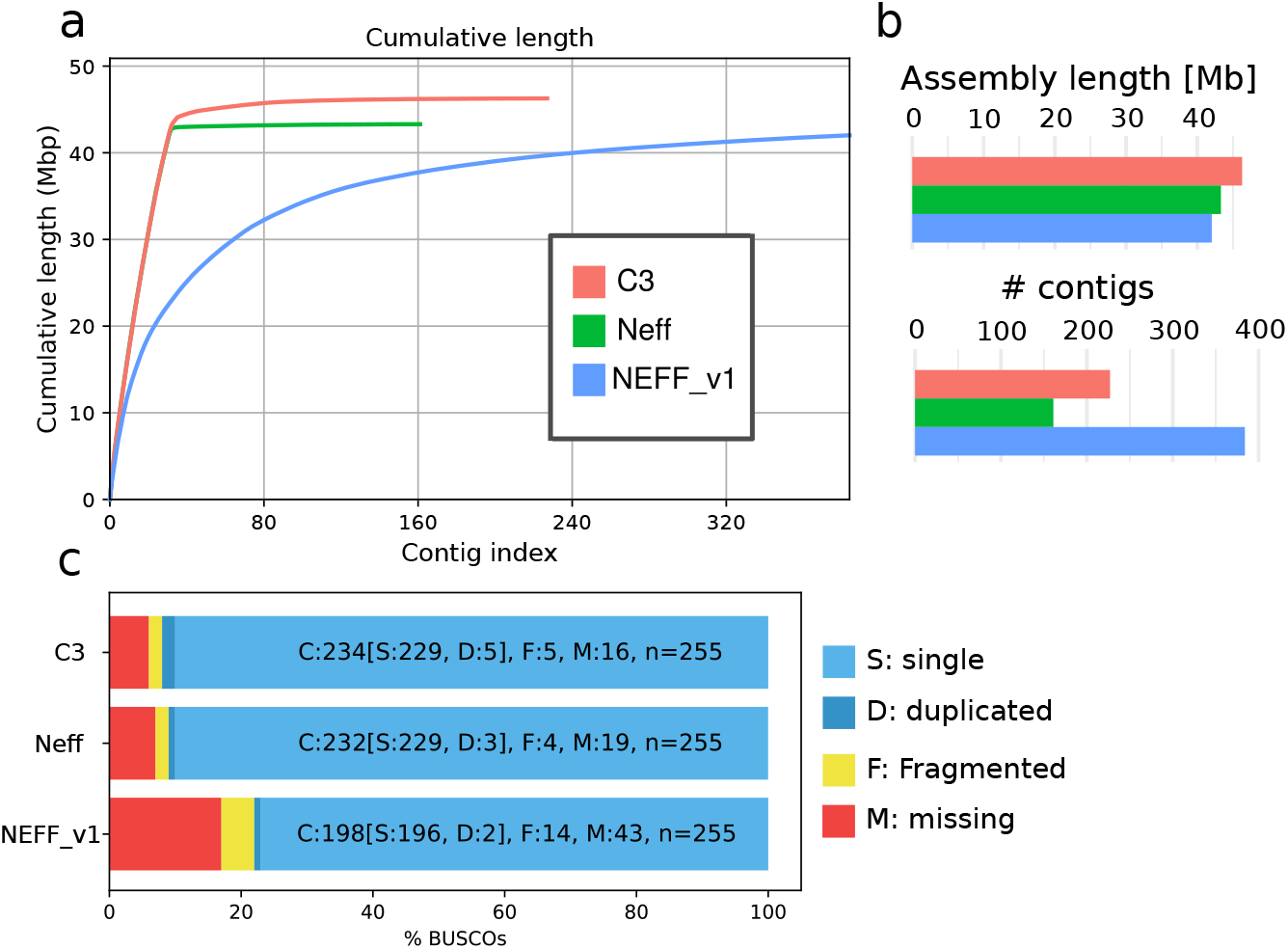
Assembly statistics for A. castellanii genomes. Comparison of genome assemblies for strains C3 and Neff, versus the previous NEFF-v1 genome assembly [25]. **a,** Cumulative length plot showing the relationship between number of contigs (largest to smallest) and length of the assembly. **b,** General continuity metrics. **c,** BUSCO statistics showing the status of universal single copy orthologs in eukaryotes for each assembly.

**Table 1.** Genome statistics for the finished assemblies of Neff, C3 (this study) and the reference Neff-v1 genome.

| Assembly | Neff-v1 | Neff | C3 |
| --- | --- | --- | --- |
| Genome size (Mbp) | 42.0 | 43.8 | 46.1 |
| # scaffolds | 384 | 111 | 174 |
| # of Ns (Mbp) | 2.6 (6.1%) | 0 (0%) | 0 (0%) |
| N50 (Mbp) | 0.3 | 1.3 | 1.4 |
| Largest scaffold (Mbp) | 2.0 | 2.5 | 2.4 |
| GC% | 57.90 | 58.44 | 58.64 |
| # protein coding genes | 14,974 | 15,497 | 16,837 |

### *A. castellanii* strains Neff and C3 have partly non-overlapping gene complements

The generation of chromosome-scale genome assemblies for two different *A. castellanii* strains afforded us the first opportunity to compare and contrast their coding capacities. We used both Broccoli [32] and OrthoFinder [33] for inference of orthologous groups. A summary of the inferred orthogroups shared by, and specific to, the Neff and C3 strains of *A. castellanii* is presented in Figure 2, with orthogroup numbers from both orthologous clustering tools included. This figure only compares Neff against C3, irrespective of orthogroup presence or absence in outgroup taxa. In this analysis, each strain-specific gene that was not assigned to an orthogroup by either program was still considered to be a single strain-specific orthogroup in order to account for the presence of genes without any orthologs across the five species. Broccoli predicted more orthogroups overall and more strain-specific genes than OrthoFinder, but predicted fewer shared orthogroups. Despite these differences, the overall trend is similar for the two outputs. The number of orthogroups shared by the two strains is roughly an order of magnitude greater than the number specific to either strain, while the C3 strain has a greater number of strain-specific orthogroups than the Neff strain as predicted by both programs.

**Figure 2.**
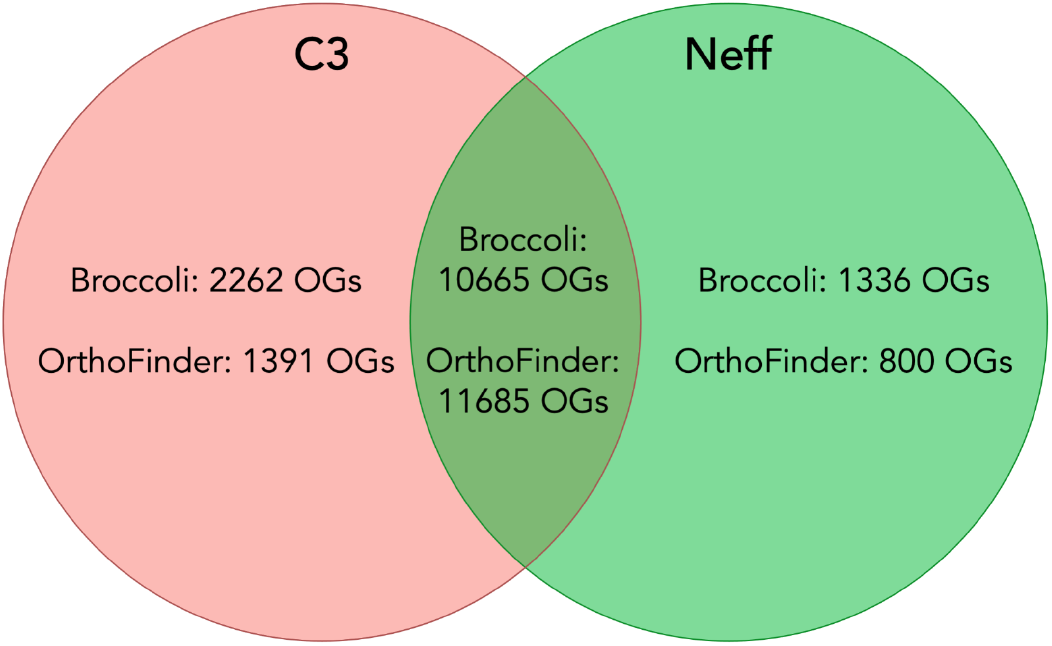
Numbers of strain-specific and shared orthologous groups in the genomes of A. castellanii strains C3 and Neff. Orthology inference was conducted with both Broccoli and OrthoFinder. Dictyostelium discoideum, Physarum polycephalum, and Vermamoeba vermiformis were used as outgroups to improve accuracy of orthogroup inference.

To investigate how similar the *A. castellanii* gene complement was to other members of Amoebozoa, *A. castellanii* orthogroups were evaluated for their presence in three outgroup species. Both Broccoli and OrthoFinder outputs were analyzed in this fashion. According to Broccoli, 43.5% of orthogroups shared by the two *A. castellanii* strains were not present in the other three amoebae, while OrthoFinder gave a figure of 48.4%. In the Neff strain, 49.1% of all orthogroups, shared or strain-specific, were not found in the three outgroup amoebae according to Broccoli, compared to 51.0% as predicted by OrthoFinder. In the C3 strain, the Broccoli results indicate that 52.4% of all orthogroups are not present in the outgroup amoebae, while 52.8% were not found in the outgroup by OrthoFinder. This is in contrast with *A. castellanii* strain C3 sharing an estimated 82.5% (Broccoli) to 89.4% (OrthoFinder) of its orthogroups with the Neff strain, and the Neff strain sharing an estimated 88.9% (Broccoli) to 93.6% (OrthoFinder) of its orthogroups with the C3 strain.

### *A. castellanii* accessory genes show strain-specific functional enrichment

In an attempt to gain insight into the functional significance of strain-specific genes in the C3 and Neff genomes, the top 30 most significantly enriched terms were identified by topGO and plotted in order of decreasing p-value for each strain/ontology combination (Supplementary Figures S8–S13). Notably, among C3-specific genes, only two terms were found to be statistically significantly enriched for each of the three ontologies at a 95% confidence level. Among Neff-specific genes, only one term was significantly enriched in each of the ‘cellular component’ and ‘molecular function’ ontologies, while three were significantly enriched in the ‘biological process’ ontology.

In C3, enriched molecular functions were ‘GTP binding’ (p = 5e-5) and ‘protein serine/threonine phosphatase activity’ (p = 0.037), enriched biological processes were ‘small GTPase mediated signal transduction’ (p = 8.5e-5) and ‘ubiquitin-dependent protein catabolic processes’ (p = 0.029), and enriched cellular components were ‘RNA polymerase II core complex’ (p = 0.026) and ‘the Golgi membrane’ (p = 0.036). In Neff, the enriched molecular function was ‘DNA helicase activity’ (p = 0.0071), enriched biological processes were ‘telomere maintenance’ (p = 0.0027), ‘protein homooligomerization’ (p = 0.0135), and ‘DNA replication’ (p = 0.0403), and the enriched cellular component was ‘virion parts’ (p = 0.012). When searched against the nr database with BLASTp [34], the Neff genes found to be responsible for both DNA helicase activity enrichment and telomere maintenance enrichment had their best BLAST hits to PIF1 5’-to-3’ DNA helicases, those responsible for protein homooligomerization enrichment had their best BLAST hits to K+ channel tetramerization domains, and the gene annotated as being a virion part had its best BLAST hits to major capsid protein from various nucleocytoplasmic large DNA viruses (NCLDVs).

### The Neff strain has a divergent mannose binding protein

One particular gene of interest encodes a mannose binding protein, which is known to be used as a receptor for cell entry by *Legionella* in some *A. castellanii* strains [35]. The MEEI 0184 strain of *A. castellanii,* an isolate from a human corneal infection, was used as a reference sequence, because it is the only strain in which the mannose binding protein is biochemically characterized [36, 37]. The orthologs from C3, Neff, and *Acanthamoeba polyphaga* were retrieved, and all four sequences were aligned (Figure S14). The percent identity of each sequence to the reference was calculated over the sites in the alignment where the *A. polyphaga* sequence was not missing (Table 2). The C3 homolog was found to be 99.5% identical to the MEEI 0184 homolog, whereas the Neff and *A. polyphaga* proteins were more divergent, sharing 91.6% and 97.2% identity to MEEI 0184, respectively. Despite being of the same species as the reference, the Neff strain homolog was found to be much more divergent than the *A. polyphaga* sequence is from the other two *A. castellanii* strains. Interestingly, we observed that *L. pneumophila* replicates worse in the Neff strain than the C3 strain in culture. This phenotype may result from impaired receptor-mediated entry by *Legionella* into Neff cells due to differences in the receptor encoding gene.

**Table 2.** Identity of mannose binding proteins from *A. polyphaga* and *A. castellanii* strains Neff and C3 to their homolog in *A. castellanii* strain MEEI 0184 across 788 sites of a 834-site amino acid alignment. The first 46 sites of the alignment were excluded from the calculation because the 5’ end of the gene in *A. polyphaga* was missing due to a truncated contig.

| Strain | Identity | Gaps |
| --- | --- | --- |
| Neff | 757/826 (91.6%) | 1/826 (0.12%) |
| C3 | 821/825 (99.5%) | 0 |
| *A. polyphaga* | 802/825 (97.5%) | 0 |

### Spatial organisation of the *A. castellanii* genome

To our knowledge, no Hi-C contact maps have been generated from species of Amoebozoa. Therefore, the Hi-C reads we used to generate the chromosome-scale scaffolding of two *A. castellanii* genomes also offer the opportunity to reveal the average genome folding in a species of this clade. Hi-C reads were realigned along the new assemblies of both the C3 and Neff strains to generate genome-wide contact maps. Visualising the Hi-C contact maps of both genomes shows that *A. castellanii* chromosomes are well resolved in our assemblies (Fig. 3). In Neff, the highest intensity contacts are concentrated on the main diagonal, suggesting an absence of large-scale mis-assemblies. On the other hand, the C3 assembly retains a few mis-assembled blocks, mostly in the rDNA region where tandem repeats could not be resolved correctly with the data available to us. However, for both strains the genome-wide contact maps reveal a grid-like pattern, with contact enrichment between chromosome extremities resulting in discrete dots. These contacts can be interpreted as a clustering of the telomeres, or subtelomeres, of the different chromosomes (Fig. 3a). Based on the presence of these inter-telomeric contacts patterns, Hi-C contact maps suggest the presence of at least 35 chromosomes in both strains, ranging from roughly 100 kbp to 2.5 Mbp in length (Fig. S15). Additionally we found 100 copies of 5S rDNA dispersed across most chromosomes for both strains, and 18S/28S rDNA genes show increased contacts with subtelomeres (Fig. 3a).

**Figure 3.**
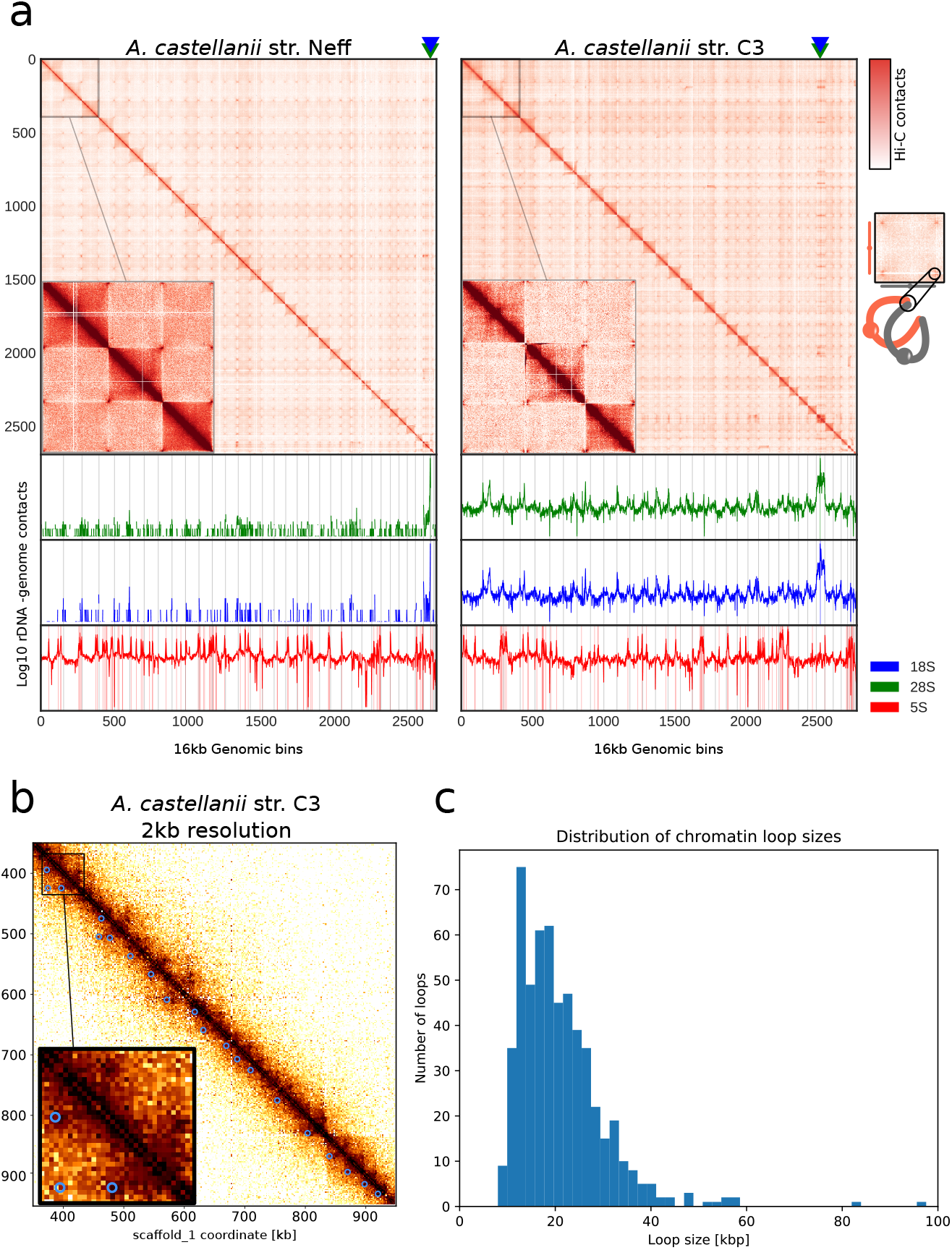
Spatial organisation of the A. castellanii genome. **a,** Top: Whole genome Hi-C contact maps of the Neff (left) and C3 (right) genomes, with a magnification of the 3 largest scaffolds. The genomes are divided into 16 kbp bins, and each pixel represents the contact intensity between a pair of bins. Each scaffold is visible as a red square on the diagonal. In both strains, there is an enrichment of inter-scaffold contacts towards telomeres, suggesting a spatial clustering of telomeres, as shown on the model in the right margin. Bottom: 4C-like representation of spatial contacts between rDNA and the rest of the genome. Scaffolds are delimited by grey vertical lines. Contacts of all rDNAs are enriched towards telomere. The genomic position of 18S and 28S genes are highlighted with triangles on the top panel and the occurences of 8S rDNA sequences are shown with vertical red lines on the bottom panel. **b,** High resolution the contact map for a region of the C3 genome showing chromatin loops detected by Chromosight as blue circles. **c,** Size distribution of chromatin loops detected in the C3 strain.

In addition to large, interchromosomal subtelomeric contacts, we also explored the existence of intrachromosomal chromatin 3D structures in the contact maps using Chromosight, a program that detects patterns reflecting chromatin structures on Hi-C contact maps [38]. For both strains, Chromosight identified arrays of chromatin loops along chromosomes, as well as boundaries separating chromatin domains (Fig. 3b). Most chromatin loops are regularly spaced, with a typical size of 20 kbp (Fig. 3c). The chromatin domains correspond to discrete squares along the diagonal (Fig. S3a). We overlapped all predicted genes in the C3 genome with the domain borders detected from Hi-C data and measured their base expression using RNA-seq we generated from that strain (see Methods). We selected the closest gene to each domain border and found that the genes overlapping domain boundaries are overall more highly expressed than those that do not (Fig. S2c). In addition, the analysis showed that gene expression is negatively correlated with the distance to the closest domain border (Fig. S2d). We performed the same comparison using chromatin loop anchors instead of domain borders. To a lesser extent, genes overlapping chromatin loops are also associated with higher expression (Fig. S2a), although it is not correlated with the distance from the closest loop (Fig. S2b). Altogether, these results suggest that the chromatin structures observed in *cis* are both associated with gene expression, although the association between gene expression and chromatin loop anchors is likely due to their co-localization with domain borders (Fig. S2e). Some microorganisms (e.g. budding yeasts and euryarchaeotes) organize their chromosomes into micro-domains that correspond to expressed genes [39, 40]. Our findings in *A. castellanii* bear an interesting similarity to this type of organization.

### *L. pneumophila* infection induces chromatin loop changes enriched in infection-related functions

The generation of near-complete assemblies allowed us to tackle the question of whether *L. pneumophila* infection impacts the 3D folding and transcription of the *A. castellanii* C3 strain genome. We harvested cultured *A. castellanii* cells before and 5 hours following infection by *L. pneumophila* strain Paris [41] (Methods). The cells were processed using Hi-C and RNA-seq (Methods), and the resulting reads aligned against the reference genome to assess changes in the genome structure and the host transcription program, respectively. RNA-seq was performed in triplicate, and Hi-C in duplicate (Methods). To measure changes in trans-chromosomal contacts, we merged the contact maps from our replicates and applied the serpentine adaptive binning method to improve the signal-to-noise ratio [42]. We then computed average interactions between each pair of chromosomes before and after infection. For each pair of chromosomes, we then used the log ratio of infected over uninfected average contacts. Following infection a global decrease in trans-subtelomeric contacts was observed, suggesting a slight de-clustering of chromosome ends (Fig. 4b). In addition, the scaffold bearing 18S and 28S rDNA (scaffold_29), as well as two other small scaffolds (35 and 36) displayed weaker interactions with other scaffolds during infection (Fig. 4a).

**Figure 4.**
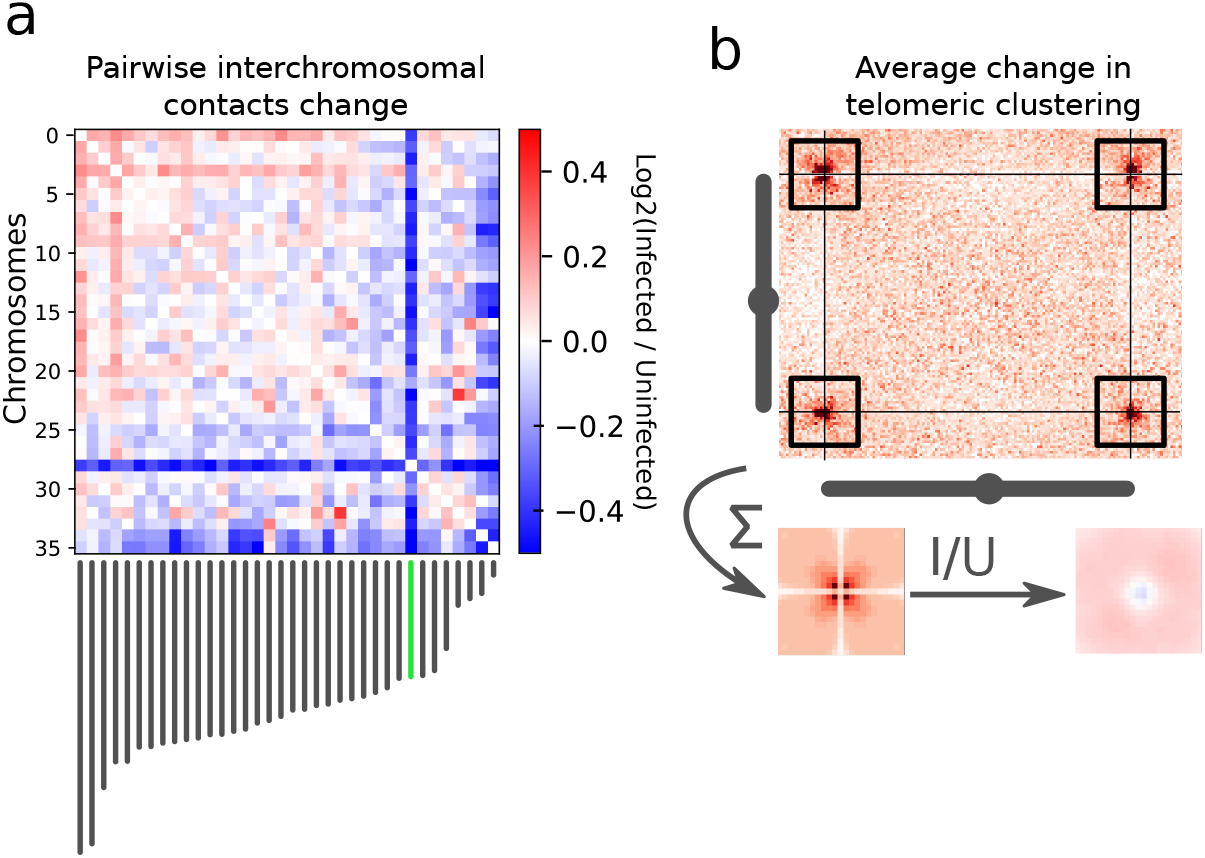
Changes in trans-chromosomal contacts between A. castellanii chromosomes following L. pneumophila infection. **a**, Average contact change during infection between each pair of chromosomes. Chromosome lengths are shown below the interaction matrix, with the chromosome bearing 18S and 28S rDNA highlighted in green **b**, Representative inter-telomeric contacts between a pair of chromosomes (Neff scaffolds 3 and 11). The average inter-telomeric contact profile generated from all pairs of chromosomes is shown as a pileup. The Log ratio between the infected (I) and uninfected (U) profiles is shown as a ratio (right).

We then assessed whether the behavior of *cis* contacts changes during infection. First, we computed the average contact frequencies according to genomic distance p(s) (Methods), which is a convenient way to unveil variations in the compaction state of chromatin [43]. The p(s) curves show a global increase in long range contacts following infection (Fig. S4b). The strengths of chromatin loops and domain borders before and 5h after infection were quantified using Chromosight [38]. However, no significant average increase or decrease in the intensity of these structures (Fig. S4a) was identified when computed over the whole genome. To focus on infection-dependent chromatin structures, we filtered the detected patterns to retain those showing the top 20% strongest change in Chromosight score during infection (either appearing or disappearing). We performed a GO term enrichment analysis for genes associated with infection-dependent chromatin loops (Methods). A significant enrichment for Rho GTPase and phosphorelay signal transduction, protein catabolism and GPI biosynthesis was found (Fig. S6a). The strongest loop changes were associated with genes encoding Rho GTPase, GOLD and SET domains as well as genes for proteins containing leucine-rich repeats and ankyrin repeats (Fig. S7).

We followed the same procedure for domain borders and found that genes associated with infection-dependent domain borders were significantly enriched in ‘amino acid transport’, ‘cyclic nucleotide biosynthetic process’, ‘protein modification’ and ‘deubiquitination’ (Fig. S6b). Our results suggest that domain borders are generally associated with highly transcribed metabolic genes, consistent with previous findings showing that such borders are associated with high transcription [44].

By analyzing the *A. castellanii* RNA-seq data after infection with *L. pneumophila*, we revealed that the expression of genes was globally impacted at 5h post infection compared to uninfected cells (Fig. S5a). This is consistent with recent results showing that transcription is globally disrupted in *A. castellanii* Neff following infection by *L. pneumophila* [22]. To investigate the relationship between this change in gene expression and chromatin structure, we assigned the closest domain border to each gene and compared their expression and border score changes during infection. For the majority of genes, we found border intensity not to be correlated with gene expression changes (Fig. S5b). Only genes undergoing extreme expression changes during infection corresponded to changes in associated borders (Fig. S5c). This raises the possibility that insulation domains in *A. castellanii* chromosomes do not dictate gene expression programs as they do in mammals.

Recently, Li et al. [22] investigated gene expression changes at 3, 8, 16 and 24h after infection of *A. castellanii* Neff by *L. pneumophila*. To further validate our finding that chromatin domains are not units of regulation in *A. castellanii*, we used these expression results and migrated the gene annotations to our C3 assembly using liftoff [45]. This allowed us to compute co-expression between gene pairs during infection (i.e., expression correlation). We found that gene pairs within the same chromatin domain did not have a higher co-expression than gene pairs from different domains at similar genomic distances (Fig. S3d).

## Discussion

### Chromosome-level assembly uncovers *A. castellanii* genome organization

Generation, analysis and comparison of the genome sequences of two *A. castellanii* strains revealed heterogeneous coverage across scaffolds, which is consistent with previous findings that *A. castellanii* has a high but variable ploidy of approximately 25n [46]. Previous estimates of the *A. castellanii* Neff karyotype using pulsed-field gel electrophoresis estimated 17 to 20 unique chromosomes ranging from 250 kbp to just over 2 Mbp [28], while our estimate suggests at least 35 unique chromosomes with a similar size range of 100 kbp to 2.5 Mbp. The discrepancy between the number of bands in the electrophoretic karyotype and our estimate may result from chromosomes of similar size co-migrating on the gel, which we were able to resolve using sequence- and contact-based information.

Considering features of the nuclear biology of *A. castellanii*, such as suspected amitosis [47] and probable aneuploidy, our finding that 5S ribosomal DNA is dispersed across all chromosomes may serve to ensure a consistent copy number of 5S rDNA in daughter cells.

It was previously estimated that *A. castellanii* has 24 copies of rDNA genes per haploid genome [48]. Our data show that both strains contain 4 times as many copies as originally thought. The decrease in interchromosomal contacts with rDNA-containing scaffolds during infection may reflect an alteration in the nucleolus structure, probably caused by a global increase in translational activity. This would be consistent with the global transcription shift observed in RNA-seq under infection conditions.

At a first glance, the contact maps show a clustering of subtelomeric regions, but do not display a Rabl conformation, where centromeres cluster to the spindle-pole body [49]. However, the precise positions of centromeres would be needed to verify that they do not co-localize with subtelomeric regions.

### Changes in chromatin structure likely reflect transcriptional changes

Infection of *A. castellanii* with *L. pneumophila* induced significant changes in chromatin loops and borders. Our analyses showed an enrichment in several interesting GO terms at the sites of these infection-induced changes, many of them consistent with known biological processes induced by *L. pneumophila* in amoebae and macrophages. Several enriched terms are related to cell cycle regulation, including mitotic cell cycle, cell cycle processes and cell cycle checkpoints (Fig. S6), which is consistent with recent results showing that *L. pneumophila* prevents proliferation of its natural host *A. castellanii* [50, 22]. *L. pneumophila*-induced alterations of the host cell cycle may serve to avoid cell cycle phases that restrict bacterial replication [51], or to prevent amoebal proliferation, which has been proposed to increase the feeding efficiency of individual amoebae [52].

Several other GO terms that we found to be enriched at infection-dependent loops or borders are related to host cell organelles, such as organelle assembly, microtubule cytoskeleton organization, protein localization to endoplasmic reticulum, mitochondrion organization, electron transport chain, or mitochondrial respiratory chain complexes (Fig. S6). This is interesting given that it is well known that during infection, *L. pneumophila* hijacks host organelles such as the cytoskeleton, the endoplasmic reticulum, and mitochondria in both amoebae and macrophages [53, 54, 55]. Indeed, mitochondrial respiration and electron transport chain complexes were recently shown to be altered in macrophages during *L. pneumophila* infection [54, 56].

Sites of infection-dependent chromatin reorganization also show enrichment in functions related to changes in the general metabolism of the host, such as biosynthetic and catabolic processes, including nucleotide and nucleoside synthesis, lipid metabolism, or transport of amino acids and metal ions. To replicate intracellularly, *L. pneumophila* acquires all its nutrients from the cytoplasm of the host cell. Therefore, it is thought that bacteria-induced modulation of the host metabolism is key to establishing a successful infection [57]. In summary, many of the GO terms associated with changes in chromatin loops and borders during infection align with the known biology of *Legionella* infection, suggesting a link between chromatin organization and many of the observed changes in host cells during infection.

It was previously shown that *L. pneumophila* infection halts host cell division and is associated with a decrease of mRNA of the *A. castellanii* CDC2b gene, a putative regulator of the *A. castellanii* cell cycle [50]. The large scale 3D changes we observed in chromatin compaction (Fig. S4b) and interchromosomal contacts (Fig. 4) are reminiscent of cell cycle changes in yeast and could suggest that the bacterium stops the host’s cell cycle at a specific checkpoint.

We identified an array of regularly spaced chromatin loops in *A. castellanii* chromosomes of approximately 20 kbp in size. This is consistent with size range of chromatin loops observed in *S. cerevisiae* during the G2/M stage [58]. This similarity in terms of regularity and size suggests that chromatin loops in *A. castellanii* may serve a similar purpose of chromosome compaction for cell division as in yeast. Our finding that DNA loop anchors and domain borders overlap highly expressed genes is also concordant with observations made in yeast and other species that domain borders are preferentially located at highly expressed genes [38, 59], and could result from their role in blocking the processing SMC complexes [60], potentially to avoid interferences between cohesin activity and transcription.

Unlike previously shown in Drosophila [61], we did not find an increase in coexpression of genes sharing the same contact domain in *A. castellanii*. This suggests chromatin domains may be caused by highly transcribed genes, and do not act as units of regulation.

### *A. castellanii* accessory genes may permit environmental adaptation

Despite the substantial number of genes predicted to be strain-specific in *A. castellanii*, few functions were found to be significantly enriched in either the Neff or C3 strain set of strain-specific genes. Of these, the most biologically interesting is the enrichment of both ‘small GTPase mediated signal transduction’ and ‘GTP binding’ genes in C3. Nearly all of the genes annotated as being involved in ‘small GTPase mediated signal transduction’ biological processes are also annotated as having ‘GTP binding’ molecular functions, which is not surprising – GTP binding is an integral part of GTPase functionality. The enrichment of these two GO terms, as well as protein serine/threonine phosphatase activity enrichment, suggests that the C3 strain may have expanded its capacity for environmental sensing and associated cellular responses by expanding gene families involved in signal transduction. Given the extensive gene repertoire in *A. castellanii* dedicated to cell signalling, environmental sensing, and the cellular response [25], which is thought to help the amoeba navigate diverse habitats and identify varied prey, it seems likely that alterations of this gene repertoire in C3 may have permitted further environmental adaptations.

Another enrichment of note is that of ‘virion parts’ in the Neff strain of *A. castellanii*. This enrichment comprises a single gene with a best BLAST hit to major capsid proteins in various NCLDVs, including a very strong hit to *Mollivirus siber-icum.* Many NCLDVs, including *Mollivirus,* are known viruses of *Acanthamoeba* spp. [62]. Although no phylogenetic analyses were performed to investigate the origin of this major capsid protein gene in the Neff genome, it seems plausible that it was acquired by lateral gene transfer during an NCLDV infection, perhaps by *Mollivirus* or some closely related virus.

The remaining enriched functions have no obvious biological significance. They could well be non-adaptive, having been generated through gene duplication, differential loss in the other surveyed amoebae, or lateral gene transfer, without conferring any notable selective advantage. An improved understanding of *Acanthamoeba* cell and molecular biology is needed to make sense of the gene enrichment data presented herein.

### Substitutions in the Neff mannose binding protein may inhibit *Legionella* entry

Alignment of the three *A. castellanii* mannose binding proteins (MBPs) and the *A. polyphaga* homolog may help explain the difference in susceptibility to *Legionella* infection between the Neff and C3 strains. The C3 strain mannose binding protein is highly similar to its counterpart in strain MEEI 0184, which was first to be biochemically characterized. The Neff strain MBP, however, is markedly more divergent than even the *A. polyphaga* MBP, which is not known to participate in *Acanthamoeba-Legionella* interactions [63]. These results are consistent with the hypothesis that the Neff strain of *A. castellanii* is not a very good host for infection by *Legionella* due to an accumulation of amino acid substitutions in its mannose binding protein, substitutions that may prevent *Legionella* from binding to this protein during cell entry. Whether or not *A. castellanii* uses its MBP for feeding or recognition of potential pathogens like *Legionella* is at present unclear, but it is worth noting that the Neff strain has been in axenic culture since 1957, so it may be that relaxed selective pressure on this protein, combined with repeated population bottlenecking during culture maintenance, has allowed for mutations in the Neff strain MBP gene to accumulate. At the present time, without available genome data for strains more closely related to the Neff strain, it cannot be determined whether these mutations arose in nature or in culture. However, given that the divergence of the *A. polyphaga* ortholog to the MEEI 0184 strain is much less than that of the Neff strain, despite all four strains having similar lifestyles in nature, evolution of the Neff strain since being deposited in the culture collection seems likely.

## Methods

### Strains and growth conditions

*A. castellanii* strains Neff and C3 were grown on amoeba culture medium (2% Bacto Tryptone, 0.1% sodium citrate, 0.1% yeast extract), supplemented with 0.1 M glucose, 0.1 mM CaCl_2_, 2.5 mM KH_2_PO_4_, 4 mM MgSO_4_, 2.5 mM Na_2_HPO_4_, 0.05 mM Fe_4_O_21_P_5_ at 20°C. *L. pneumophila* strain Paris was grown for 3 days on N-(2-acetamido)-2-amino-ethanesulfonic acid (ACES)-buffered charcoal-yeast (BCYE) extract agar, at 37 °C.

### Infection timecourse

Infection of *A. castellanii* C3 with *L. pneumophila* was performed using MOI 10 over 5h in infection medium (0.5% sodium citrate supplemented with 0.1 mM CaCl_2_, 2.5 mM KH_2_PO_4_, 4 mM MgSO_4_, 2.5 mM Na_2_HPO_4_, 0.05 mM Fe_4_O_21_P_6_ at 20°C. At 5h post-infection, amoebae were collected in a 15 mL tube, pelleted by centrifugation at 300 *g* for 10 minutes and washed twice in PBS, then crosslinked in 3% formaldehyde during 20 minutes at room temperature (RT) with gentle shaking. 2.5 M glycine was then added to reach a final concentration of 0.125 M over 20 minutes, centrifuged, washed, and pellets were stored at −80°C until DNA extraction.

### Library preparations

#### Hi-C

Cell pellets were suspended in 1.2ml H2O and transferred to CK14 Precellys tubes. Cells were broken with Precellys (6 cycles: 30 sec ON / 30 sec OFF) at 7500 RPM and transferred into a tube. All Hi-C libraries for *A. castellanii* strains C3 and Neff were prepared using the Arima kit and protocol with only the *DpnII* restriction enzyme. Libraries were sequenced to produce 35 bp paired-end reads on an Illumina NextSeq machine.

#### Short-read sequencing

Illumina libraries SRX12218478 and SRX12218479 were prepared from A. castellanii strains C3 and Neff genomic DNA, respectively, and sequenced by Novogene at 2×150 bp on an Illumina Novaseq 6000 machine.

For SRX4625411, a PCR-free library was prepared and sequenced by Genome Québec from purified A. castellanii strain Neff genomic DNA. The library was barcoded and run with other samples on an Illumina HiSeq X Ten instrument, producing 150 bp paired-end reads.

#### RNA-seq

Poly-A selected libraries were prepared from purified *A. castellanii* total RNA. *A. castellanii* strain C3 RNA-seq libraries were prepared using the stranded mRNA Truseq kit from Illumina and sequenced in single-end mode at 150 bp on an Illumina NextSeq machine.

For *A. castellanii* strain Neff (SRX7813524), the library was prepared and sequenced by Génome Québec. The library was barcoded and run with other samples on an Illumina NovaSeq 6000 instrument, producing 300 bp paired-end reads.

#### Nanopore sequencing

For SRX12218489 and SRX12218490, DNA was extracted from *A. castellanii* strains Neff and C3 using the QIAGEN Blood and Cell Culture DNA Kit (Qiagen) following the specific recommendations detailed by Oxford Nanopore Technologies in the info sheet entitled “High molecular weight gDNA extraction from cell lines (2018)” in order to minimize DNA fragmentation by mechanical constraints. Nanopore libraries were prepared with the ligation sequencing kit LSKQ109, flowcell model MIN106D R9. Basecalling was performed using Guppy v2.3.1-1.

For other libraries, genomic DNA samples were obtained from *A. castellanii* strain Neff using an SDS-based lysis method, followed by digestion with RNase A, then proteinase K, and then a phenol-chloroform-based extraction. DNA samples were cleaned with QIAgen G/20 Genomic Clean-up columns using the manufacturer’s protocol, but with double the number of wash steps. Four different libraries were prepared, using the SQK-RAD003 Rapid Sequencing Kit (SRX4620962), the SQK-LSK308 1D2 Ligation Sequencing Kit (SRX4620963), the SQK-RAD004 Rapid Sequencing Kit (SRX4620964), and the SQK-LSK108 Ligation Sequencing Kit (SRX4620965). The SQK-LSK308 and SQK-RAD003 libraries were sequenced on FLO-MIN107 flow cells, and the SQK-LSK108 and SQK-RAD004 libraries were both sequenced on a FLO-MIN106 flow cell. All four libraries were basecalled with Albacore 2.1.7, as they were sequenced prior to the release of Guppy. Adapters were removed from the basecalled reads using Porechop v0.2.3.

### Genome assembly

Nanopore reads were filtered using filtlong v0.2.0 with default parameters to keep the best 80% reads according to length and quality. Illumina shotgun libraries were used as reference for the filtering. A *de novo* assembly was generated from the raw (filtered) Nanopore long reads using flye v2.3.6 with three iterations of polishing. The resulting assembly was polished using both Nanopore and Illumina reads with HyPo v1.0.1. Contigs from the polished assembly bearing more than 60% of their sequence or 51% identity to the mitochondrial sequence from the NEFF_v1 assembly were separated from the rest of the assembly to prevent inclusion of mitochondrial contigs into the nuclear genome during scaffolding. Polished nuclear contigs were scaffolded with Hi-C reads using instagraal v0.1.2 with default parameters. Instagraal-polish was then used to fix potential errors introduced by the scaffolding procedure. Mitotic contigs were then added at the end of the scaffolded assembly and the final assembly was polished with the Illumina shotgun library data using two rounds of pilon polishing. The resulting assembly was edited manually to remove spurious insertion of mitochondrial contigs in the scaffold and other contaminants. The final assembly was polished again using pilon with Rcorrector-corrected reads [64]. Minimap2 v2.17 [65] was used for all long reads alignments, and bowtie2 v2.3.4.1 for short reads alignments.

### Genome annotation

The structural genome annotation pipeline employed here was implemented similarly as described in [66]. Briefly, RNA-Seq reads were mapped to the genome assembly using STAR v2.7.3a [67], followed by both *de novo* and genome-guided transcriptome assembly by Trinity v2.12.0 [68]. Both runs of Trinity were performed with jaccard clipping to mitigate artificial transcript fusions. The resulting transcriptome assemblies were combined and aligned to the genome assembly using PASA v2.4.1 [69]. Protein sequences were aligned to the genome using Spaln v2.4.2 [70] to recover the most information from sequence similarity. The *ab initio* predictors employed were Augustus v3.3.2 [71], Snap [72], Genemark v4.33 [73], and CodingQuarry v2.0 [74]. Finally, the PASA assembly, Spaln alignments, as well as Augustus, Snap and Codingquarry gene models, were combined into a single consensus with Evidencemodeler v1.1.1 [75].

Functional annotations were added using funannotate v1.5.3. [76] Repeated sequences were masked using repeatmasker. Predicted proteins were fed to Interproscan v5.22 [77], Phobius v1.7.1 [78] and Eggnog-mapper v2.0.0 [79] were used to generate functional annotations. Ribosomal RNA genes were annotated separately using RNAmmer v1.2 [80] with HMMER 2.3.2.

As described in the Availability of data and materials section, the funannotate-based script “func_annot_from_gene_models.sh” used to add functional annotations to existing gene models is provided in the Zenodo record and on the associated github repository.

### Analysis of sequence divergence

To compute the proportion of substituted positions in aligned segments between the C3 and Neff strains, the two genomes were aligned using minimap2 with the map-ont preset and -c flag. The gap-excluded sequence divergence (mismatches / (matches + mismatches) was then computed in each primary alignment and the average of divergences (weighted by segment lengths) was computed. This is implemented in the script “04_compute_seq_divergence.py” available in the genome analysis repository listed in Availability of data and materials

### Orthogroup inference

Orthogroups were inferred using the predicted proteomes of both the Neff and C3 strains, with *Dictyostelium discoideum*, *Physarum polycephalum*, and *Vermamoeba vermiformis* as outgroups to improve the accuracy of orthogroup inference. The outgroup predicted proteomes were retrieved from PhyloFisher [81]. Both Broccoli [32] and OrthoFinder [33] were run with default settings for orthogroup inference.

### Gene content comparison of Neff and C3 strains

Custom Python scripts were used to retrieve genes unique to each *A. castellanii* strain, as well as orthogroups that were shared between the two strains. Genes were only determined to be strain-specific or shared if both Broccoli and OrthoFinder assigned them as such; genes were excluded from the analysis if both tools did not agree. For both strains, functional assignments for each gene ID were extracted from funannotate output and tabulated. The tabulated assignments and strain-specific gene IDs were fed into the R package topGO [82] to analyze GO term enrichment in the strain-specific genes. Fisher’s exact test with the weight algorithm was implemented in topGO for the Neff-and C3-specific genes for each of the three ontologies (biological process, cellular component, and molecular function). When building the GOdata objects for these three ontologies, nodeSize was set to 10 for both the biological process and molecular function ontologies, and 5 for the cellular component ontology due to the lower number of GO terms in this ontology.

### Mannose Binding Protein Comparison

Mannose binding protein (MBP) amino acid sequences from three strains of *A. castellanii* (Neff, C3, and MEEI 0184) and one strain of *Acanthamoeba polyphaga* were retrieved, aligned using MAFFT-linsi (v7.475) [83], and visualized in Jalview (v2.11.1.3) [84]. The MEEI 0184 strain sequence was retrieved from NCBI (Accession: AAT37865.1), and the Neff and C3 sequences were retrieved from the predicted proteomes generated in this study with the MEEI 0184 sequence as a BLASTp [34] query. The *A. polyphaga* genome does not have a publicly available predicted proteome, so its MBP protein sequence was manually extracted from several contigs in the genome sequence (NCBI accession: GCA_000826345.1) using tBLASTn with the MEEI 0184 sequence as a query (the sequence encoding the first 8 amino acids of the protein could not be found in the genome due to a truncated contig).

### Hi-C analyses

Reads were aligned with bowtie2 v2.4.1, and Hi-C matrices were generated using hicstuff v3.0.1 [85]. For all comparative analyses, matrices were downsampled to the same number of contacts using cooltools (https://www.github.com/mirnylab/cooltools) and balancing normalization was performed using the ICE algorithm [86]. Loops and domain borders were detected using Chromosight v1.6.1 [38] using the merged replicates at a resolution of 2 kbp. We measured the intensity changes in Chromosight scores during infection using pareidolia (v0.6.1) [87] on 3 pseudo replicates generated by sampling the merged contact maps, as described in [88]. This was done to account for contact coverage heterogeneity across replicates. The 20% threshold used to select differential patterns amounts to 1.2% false detections for loops and 2.3% for borders when comparing pseudo-replicates from the same condition.

## Acknowledgements

We thank Axel Cournac, Laura Gomez Valero, Christophe Rusniok and Lyam Baudry for their comments on the bioinformatics analysis, Tobias Sahr for RNAseq library construction, Pierrick Moreau for his help with the optimization of the Hi-C protocol, Charlotte Cockram for her help with Nanopore sequencing, as well as all members of the Koszul lab and Buchrieser lab for stimulating discussions.

## Availability of data and materials

Sequencing datasets have been deposited in SRA under bioprojects PRJNA599339 and PRJNA487265. All processed data, as well as the assemblies and annotations used in this work are available on zenodo record https://zenodo.org/record/5507417. Strains supporting the findings of this study are available from the corresponding authors.

The analyses are packaged into the following snakemake pipelines available on github. Hybrid genome assembly: https://github.com/cmdoret/Acastellanii_hybrid_assembly, functional annotation of *A. castellanii:* https://github.com/cmdoret/Acastellanii_genome_annotation, analyses of genomic features in A. *castellanii*: https://github.com/cmdoret/Acastellanii_genome_analysis, changes during infection by Legionella: https://github.com/cmdoret/Acastellanii_legionella_infection.

The dataset(s) supporting the conclusions of this article are available in the Zenodo repository https://zenodo.org/record/5507417.

## Consent for publication

All authors gave their consent for publication of this manuscript.

## Authors’ contributions

C.M.D and M.J.C performed analyses. P.E. Performed infection experiments and DNA extractions. M.J.C performed DNA extraction. M.J.C and C.M.D did the Nanopore sequencing. A.T. constructed Hi-C and shotgun libraries.

M.J.C., C.M.D., B.C., M.S., M.W.G. and B.F.L. contributed to the curation and improvement of the genome assembly. All authors contributed to writing the manuscript.

## Supplementary figures

**Figure S1.**
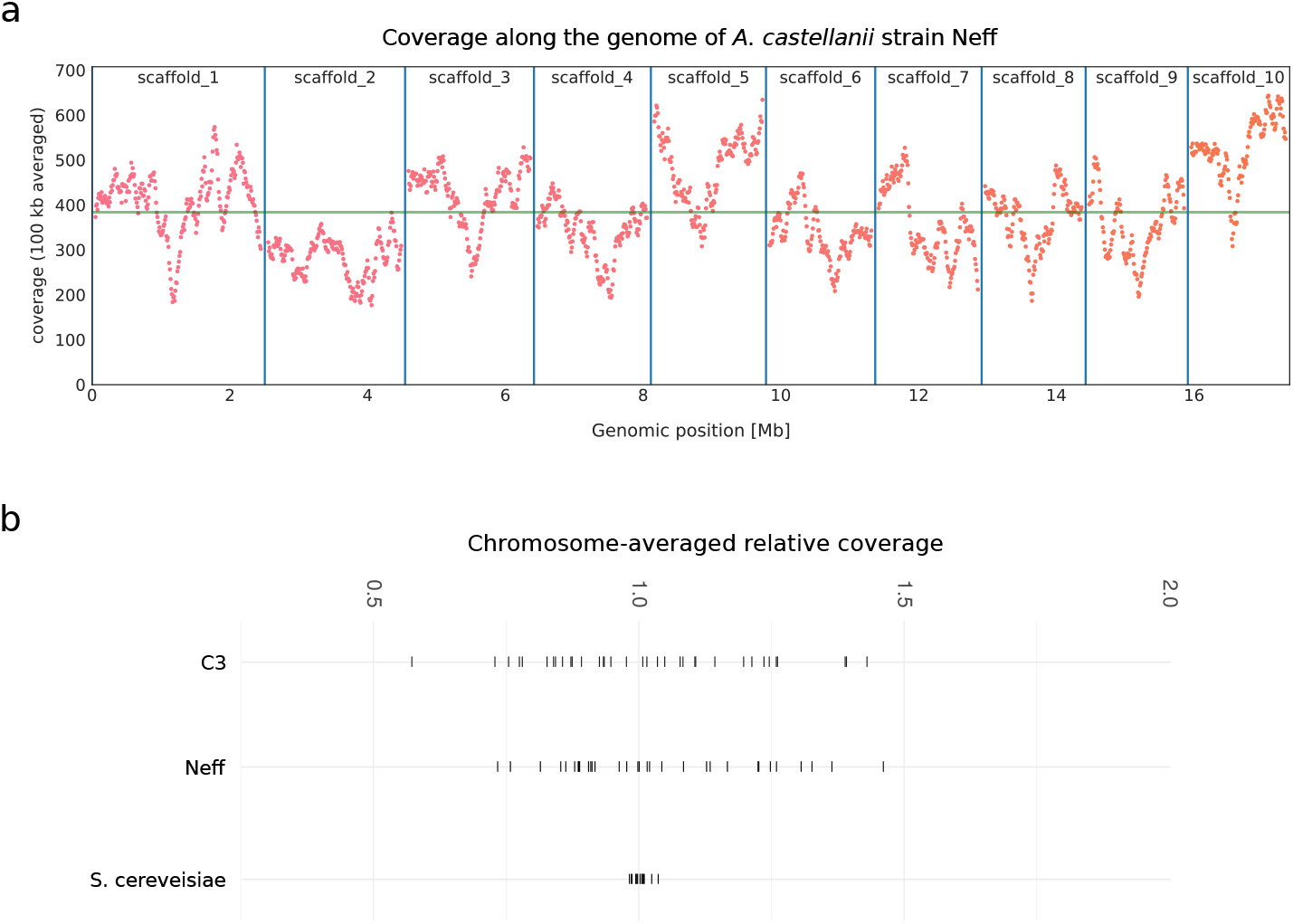
Coverage across scaffolds of A. castellanii compared to a known haploid. **a,** Illumina short-reads coverage along the 10 largest scaffolds of *A. castellanii* Neff in a 100 kbp sliding window, with the horizontal green line showing genome median coverage. **b,** Variability of median coverage per chromosome (relative to genome median) for *A. castellanii* strains C3 and Neff, and asynchronous *Saccharomyces cerevisiae* strain BY4741, a known haploid. For *S. cerevisiae*, library SRR1569870 was used.

**Figure S2.**
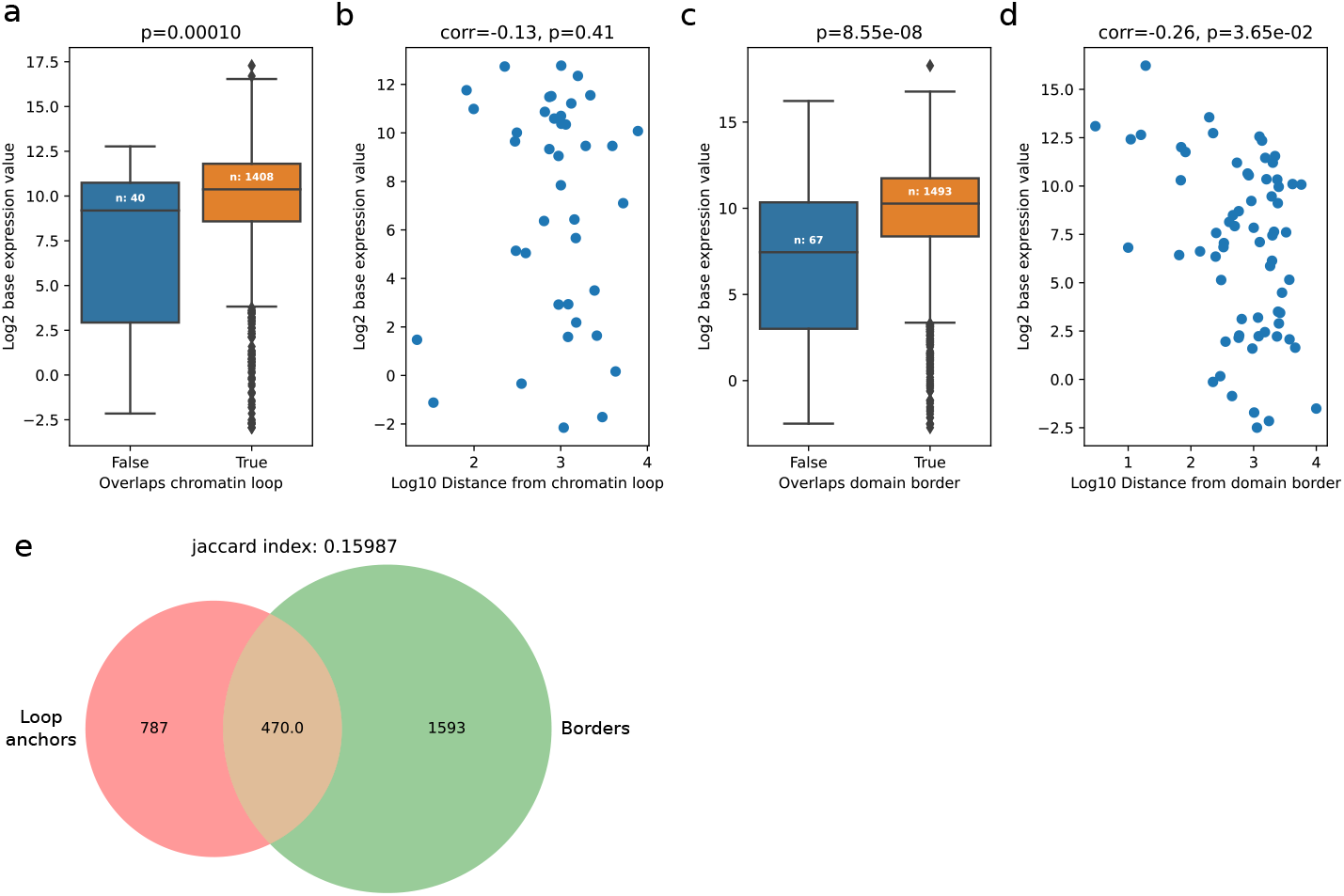
Gene expression according to position relative to chromatin structures. Expression of the closest gene to each loop anchors versus **a,** overlap status with chromatin loops and **b,** distance to closest loop. Expression of the closest gene to domain borders versus **c,** overlap status with domain borders and **d,** distance to closest border. P-values reported for overlap comparisons are obtained using Mann-Whitney U test, correlation coefficients and associated p-values are computed using Spearman’s correlation test. **e,** Overlap between chromatin loop anchors and domain borders represented as a Venn diagram.

**Figure S3.**
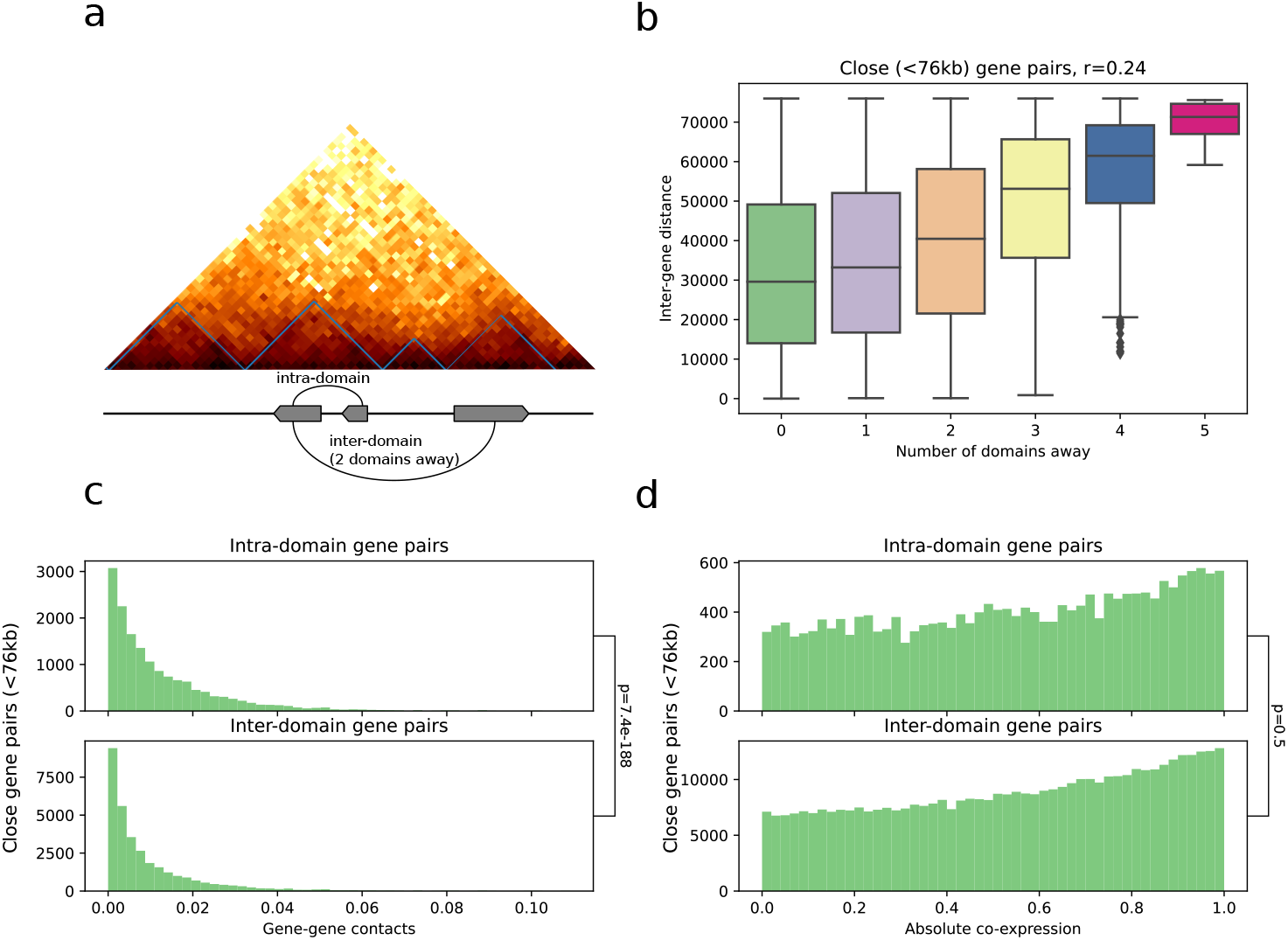
Relationship between genes and insulation domains. **a,** Example domains detected by Chromosight in the C3 strain, with theoretical genes for demonstration. **b,** Relationship between inter-gene distance and number of domains separating them. **c,** Distribution of mean inter-gene contacts according to domain separation status. **d,** Distribution of gene-pairs co-expression according to domain separation status. For all panels, only gene pairs separated by less than the median domain size were selected.

**Figure S4.**
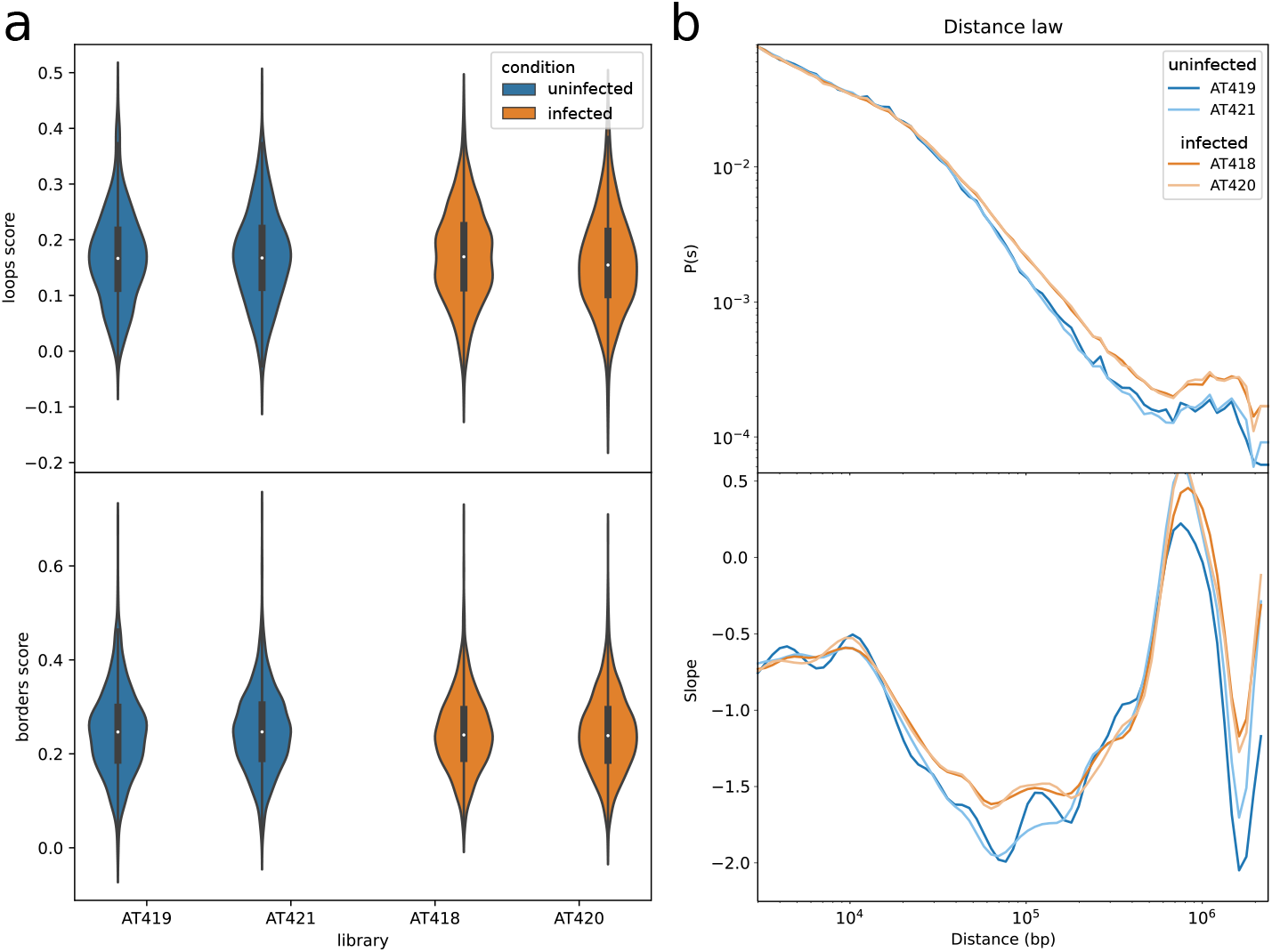
Global comparisons of infection Hi-C results between replicates. **a,** Distribution of Chromosight loops and borders scores for all 4 samples. **b,** Distance-contact decay function (denoted P(s)) and its slope.

**Figure S5.**
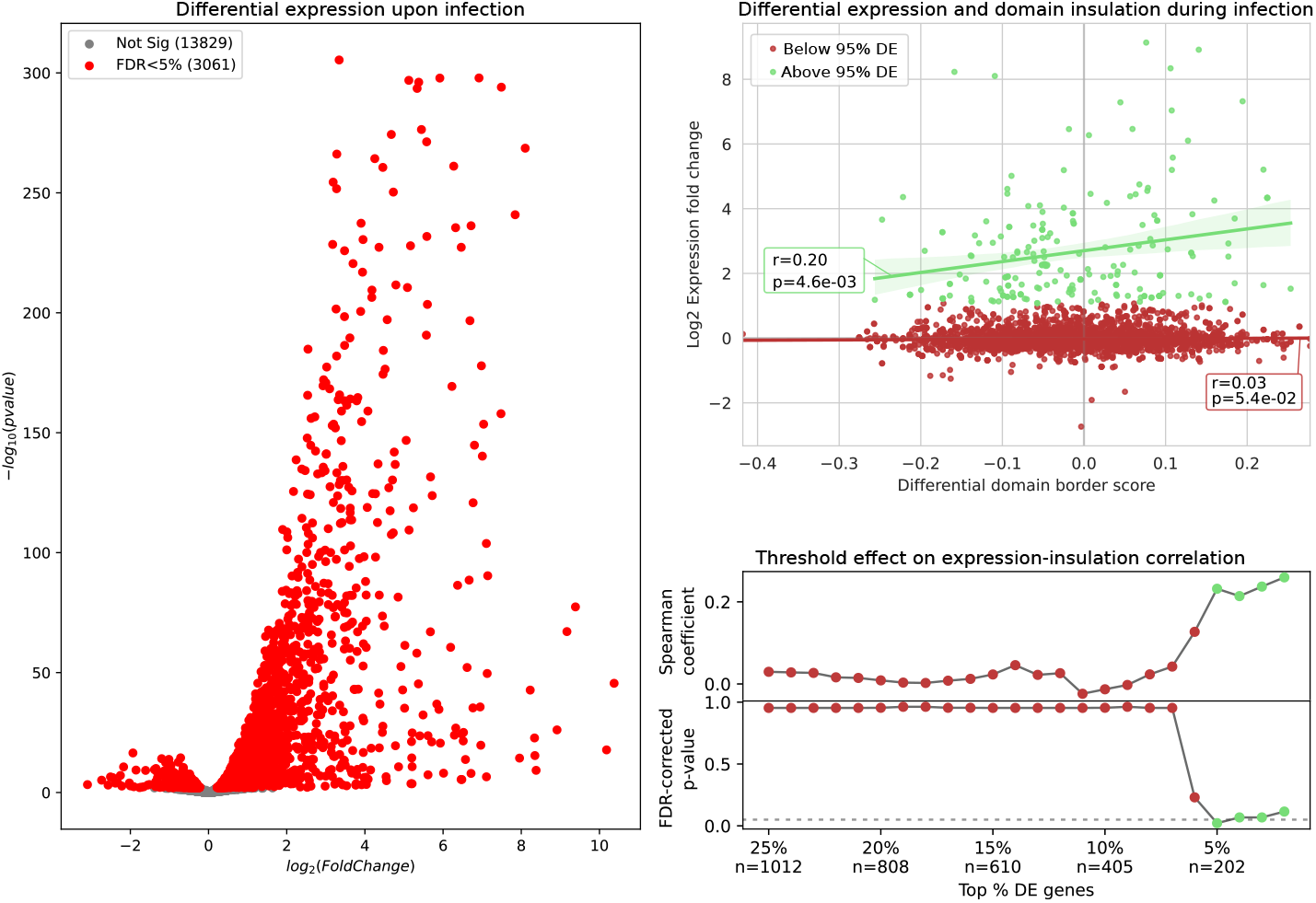
Relationship between differential expression and domain insulation during infection. **a,** Volcano plot showing differential gene expression (DE) of infected (5h p.i.) versus uninfected amoeba. Genes with significant corrected p-values (FDRj5%) are shown in red. **b,** Changes in gene expression and insulation strength of closest domain border during infection. Linear regression lines, Spearman correlation coefficients and associated p-values are shown separately for genes with extreme fold change values (95% quantile) and the rest. **c,** Spearman correlation coefficient between expression fold change and domain insulation change, and associated FDR-corrected p-values (FDRj5%) for different subsets of genes according to the threshold of extreme fold change. Values are colored according to the 95% threshold selected in b.

**Figure S6.**
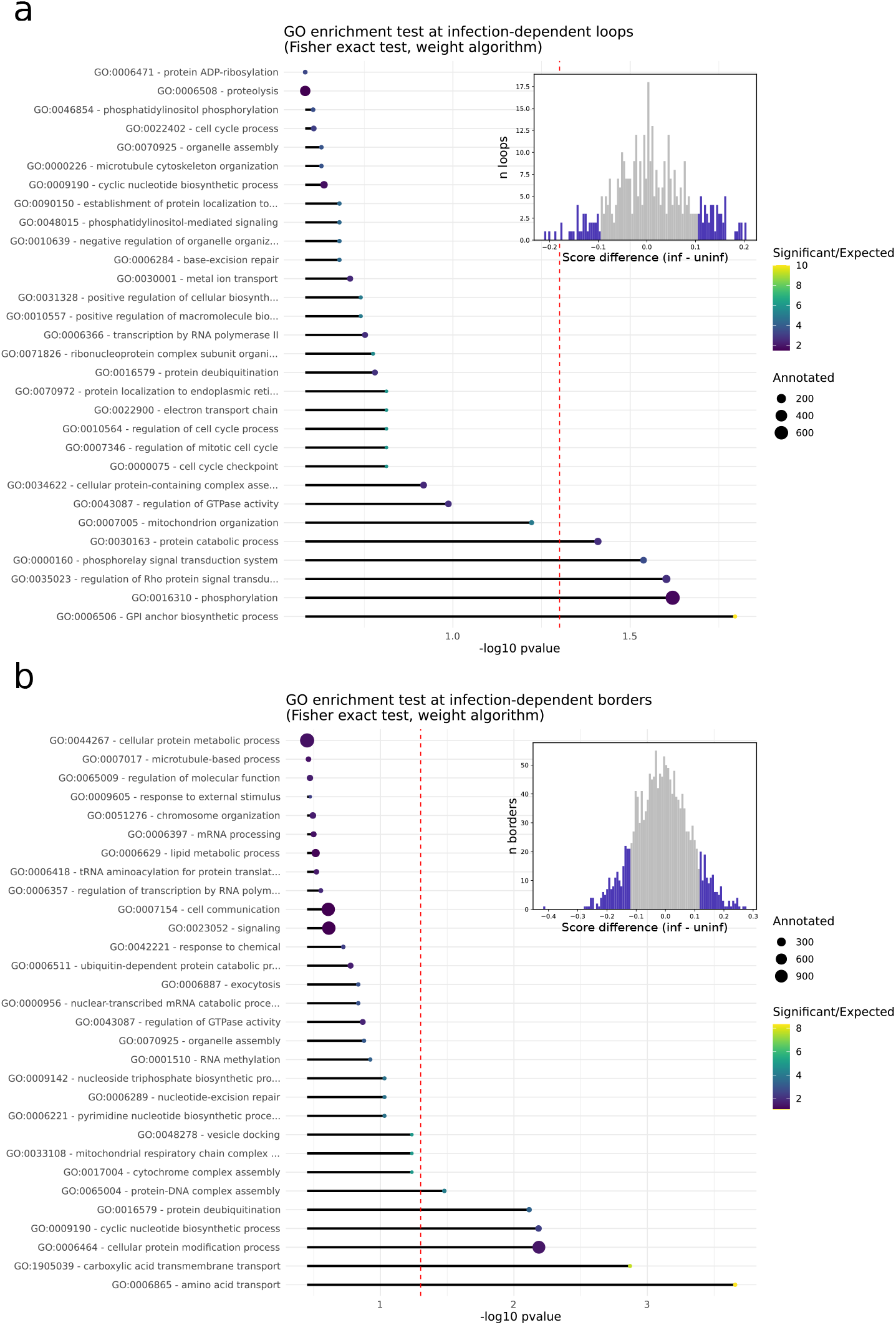
GO term enrichment test results for genes overlapping infection-dependent **a,** chromatin loops and **b,** domain borders. Histograms show the distribution of loop and border score changes during infection, with highlighted portions showing the 80% percentile threshold used to include genes in the GO enrichment test.

**Figure S7.**
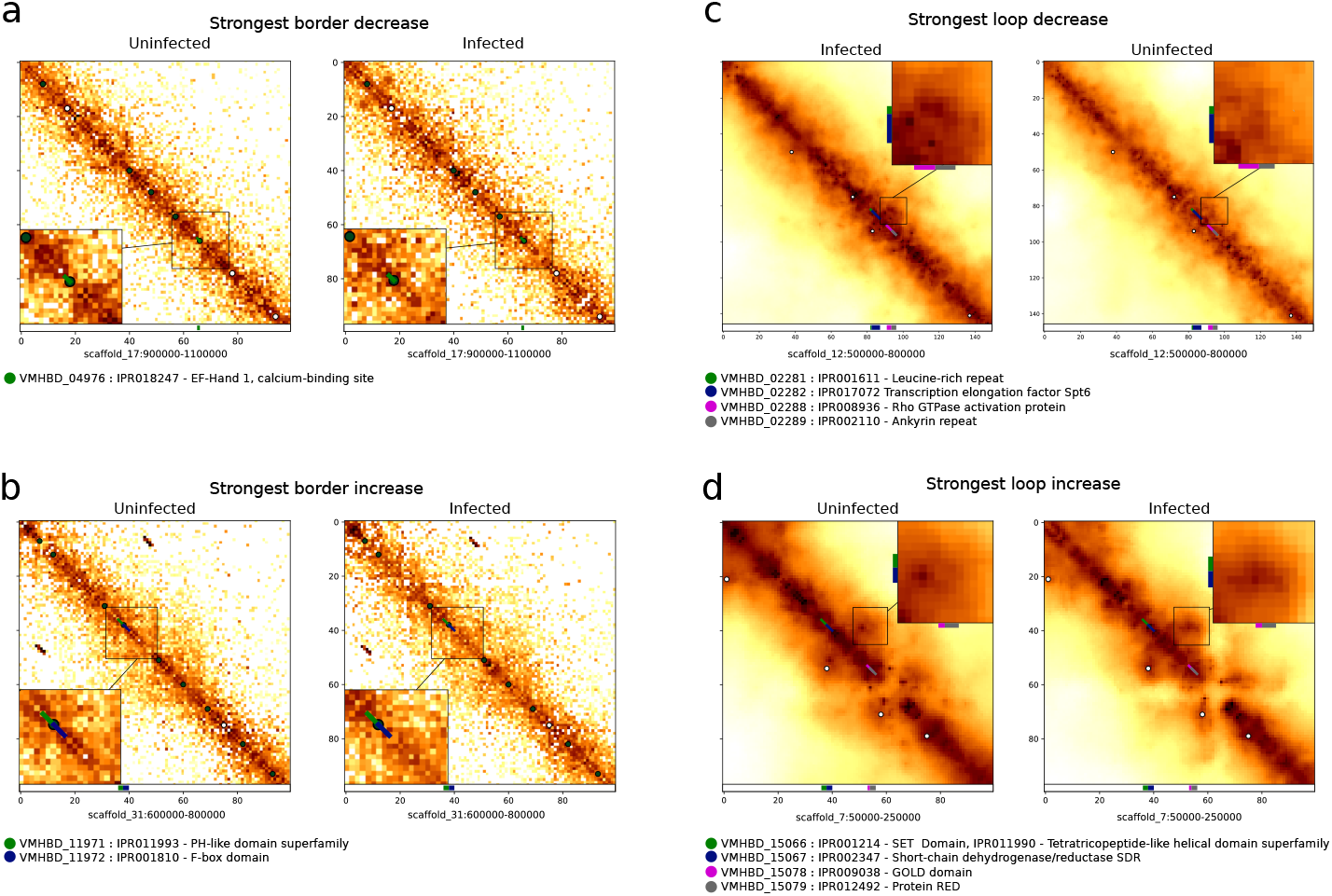
Hi-C zooms on strongest pattern changes during infection. Description of the closest genes are shown below each zoom. Balanced contact map zooms showing **a,** strongest border decrease and **b,** decrease. Serpentine-binned contact maps showing **c,** strongest loop decrease and **d,** increase.

**Figure S8.**
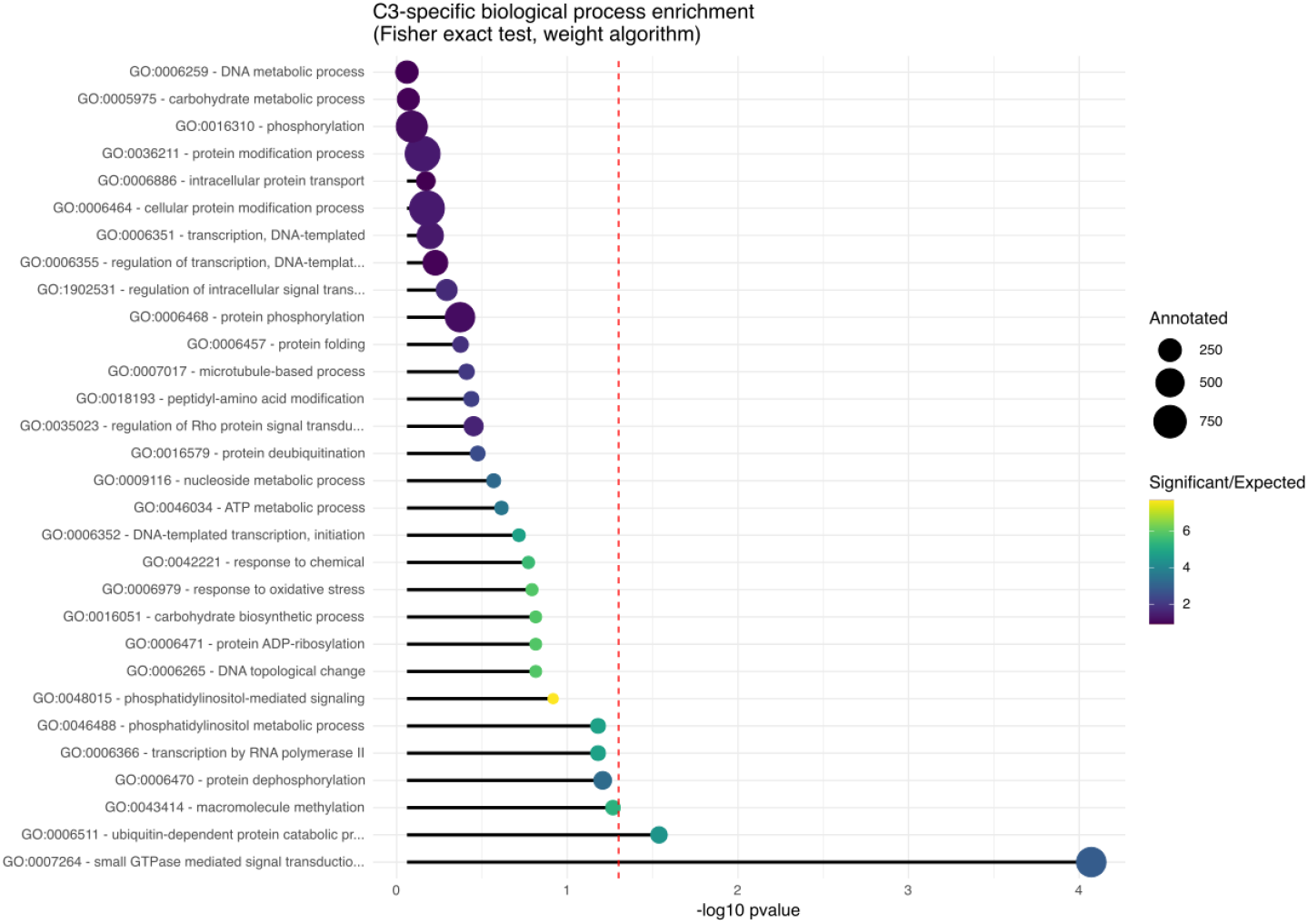
Most significant biological process GO term enrichments in genes specific to Acanthamoeba castellanii strain C3. Enrichment was determined using topGO, with nodeSize set to 10 when building the GOdata object. The size of the circles at the end of the bars represents the number of genes annotated under that GO term in the genome, and the colour scale of the circles represents the ratio of how many genes were found in the strain-specific set for that term compared to how many were expected.

**Figure S9.**
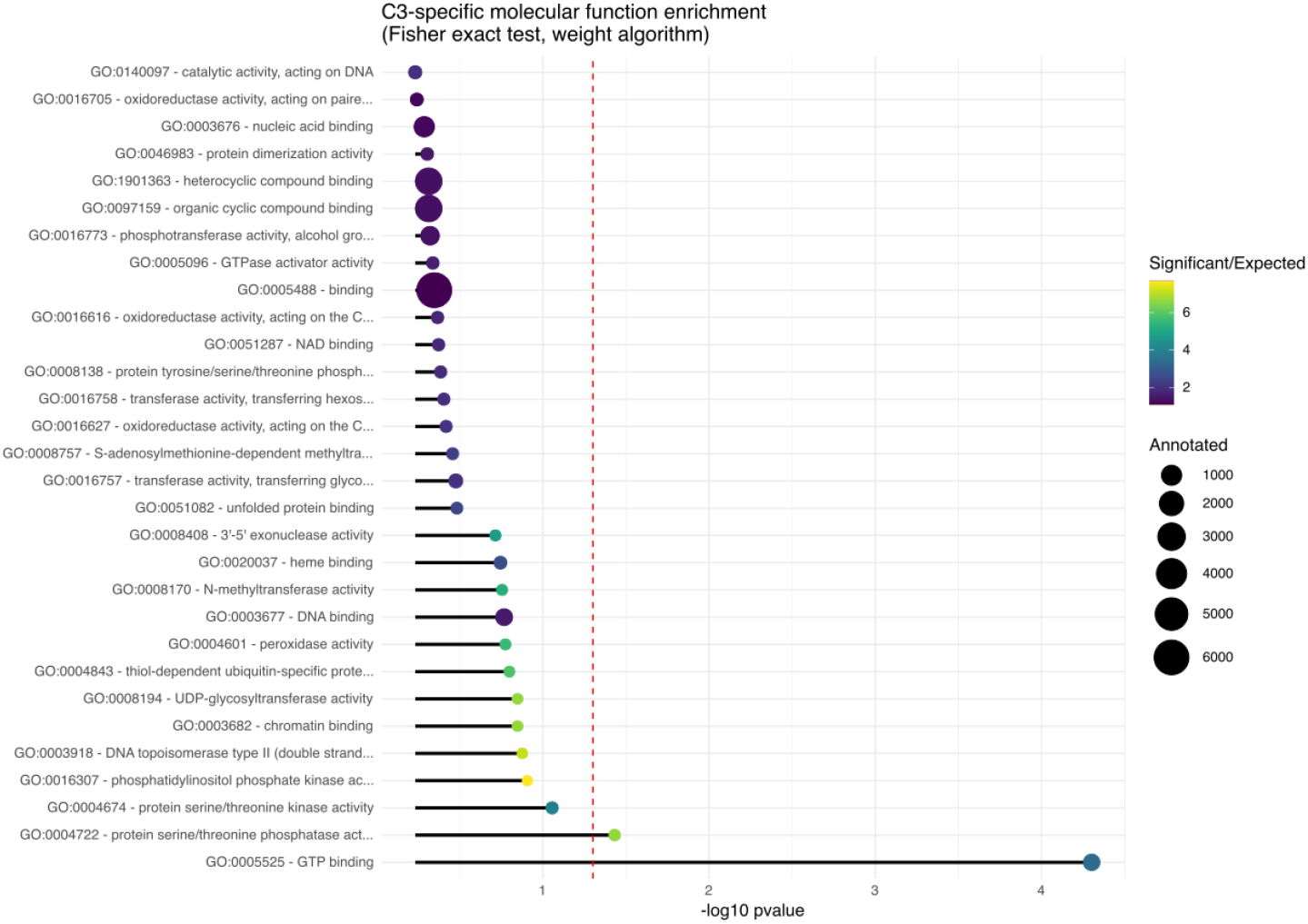
Most significant molecular function GO term enrichments in genes specific to Acanthamoeba castellanii strain C3. Enrichment was determined using topGO, with nodeSize set to 10 when building the GOdata object. The size of the circles at the end of the bars represents the number of genes annotated under that GO term in the genome, and the colour scale of the circles represents the ratio of how many genes were found in the strain-specific set for that term compared to how many were expected.

**Figure S10.**
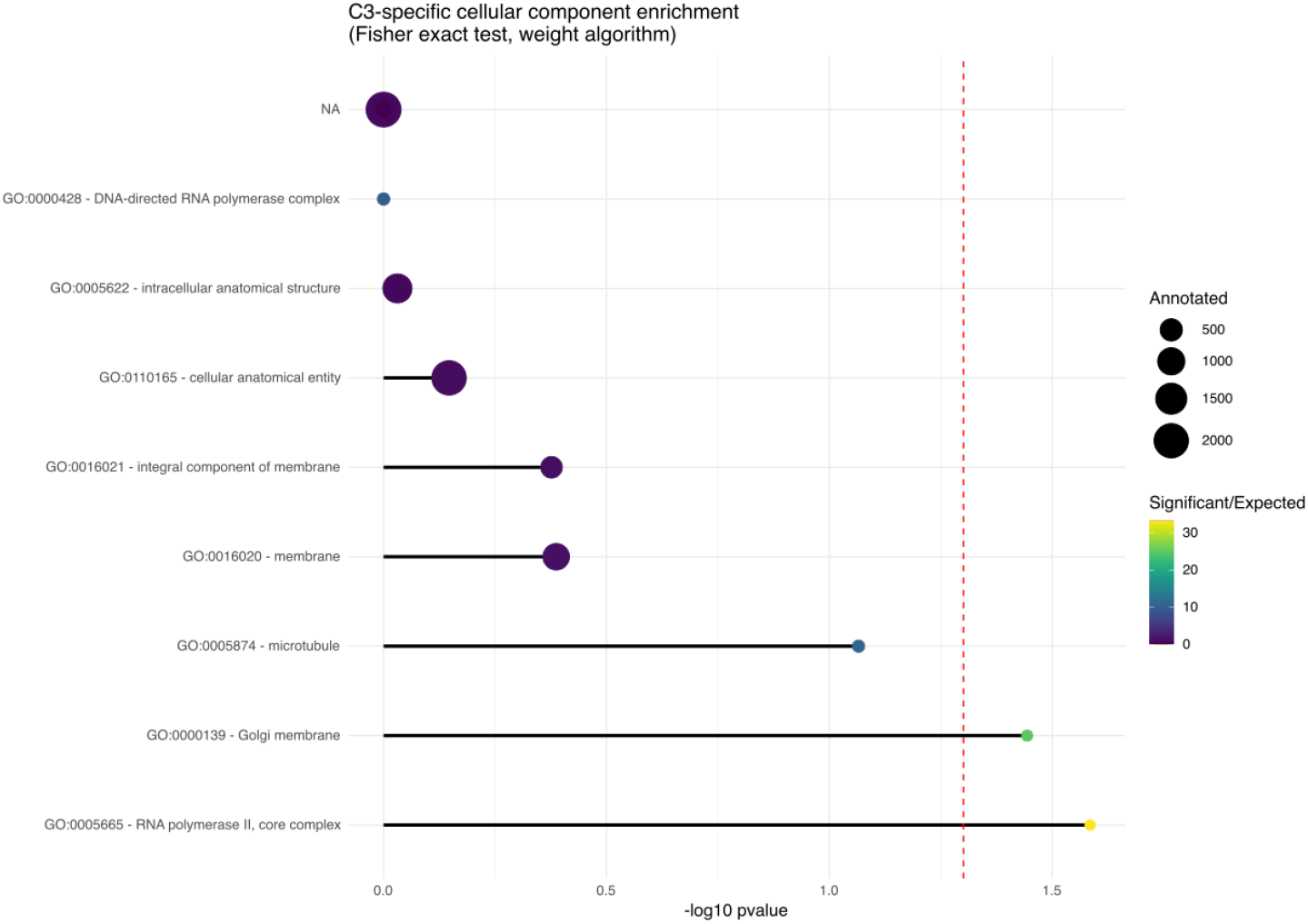
Most significant cellular component GO term enrichments in genes specific to Acanthamoeba castellanii strain C3. Enrichment was determined using topGO, with nodeSize set to 5 when building the GOdata object. The size of the circles at the end of the bars represents the number of genes annotated under that GO term in the genome, and the colour scale of the circles represents the ratio of how many genes were found in the strain-specific set for that term compared to how many were expected.

**Figure S11.**
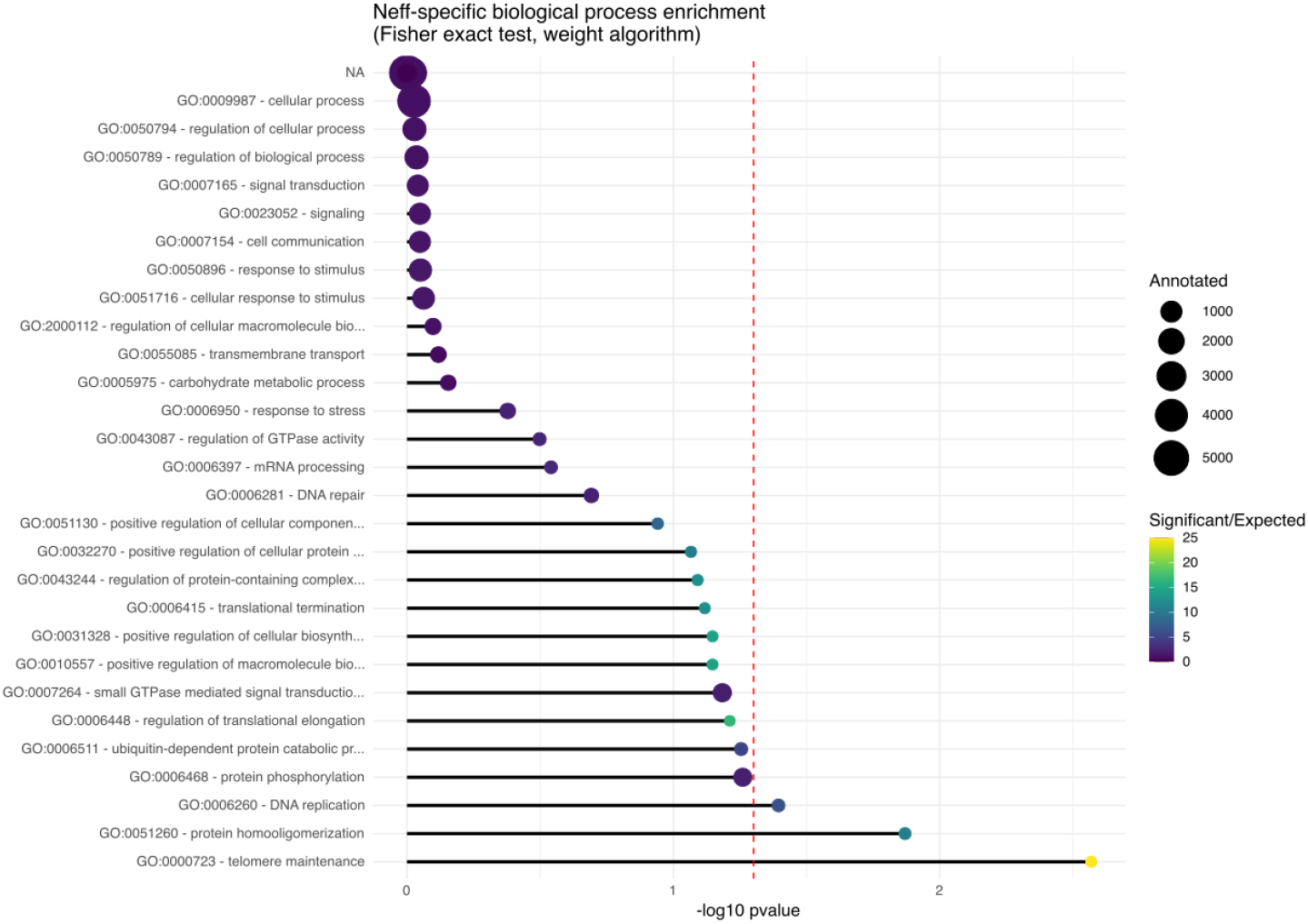
Most significant biological process GO term enrichments in genes specific to Acanthamoeba castellanii strain Neff. Enrichment was determined using topGO, with nodeSize set to 10 when building the GOdata object. The size of the circles at the end of the bars represents the number of genes annotated under that GO term in the genome, and the colour scale of the circles represents the ratio of how many genes were found in the strain-specific set for that term compared to how many were expected.

**Figure S12.**
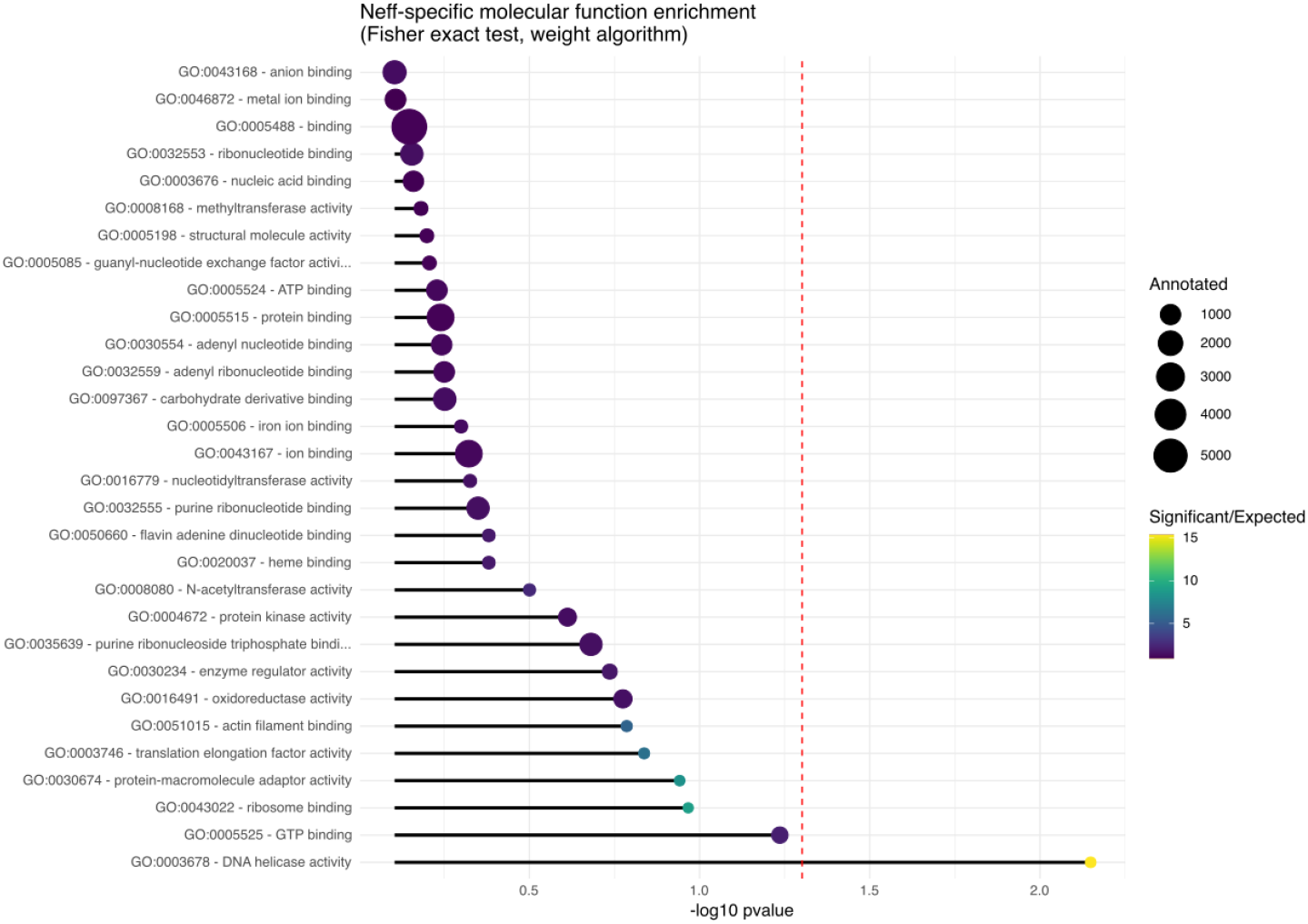
Most significant molecular function GO term enrichments in genes specific to Acanthamoeba castellanii strain Neff. Enrichment was determined using topGO, with nodeSize set to 10 when building the GOdata object. The size of the circles at the end of the bars represents the number of genes annotated under that GO term in the genome, and the colour scale of the circles represents the ratio of how many genes were found in the strain-specific set for that term compared to how many were expected.

**Figure S13.**
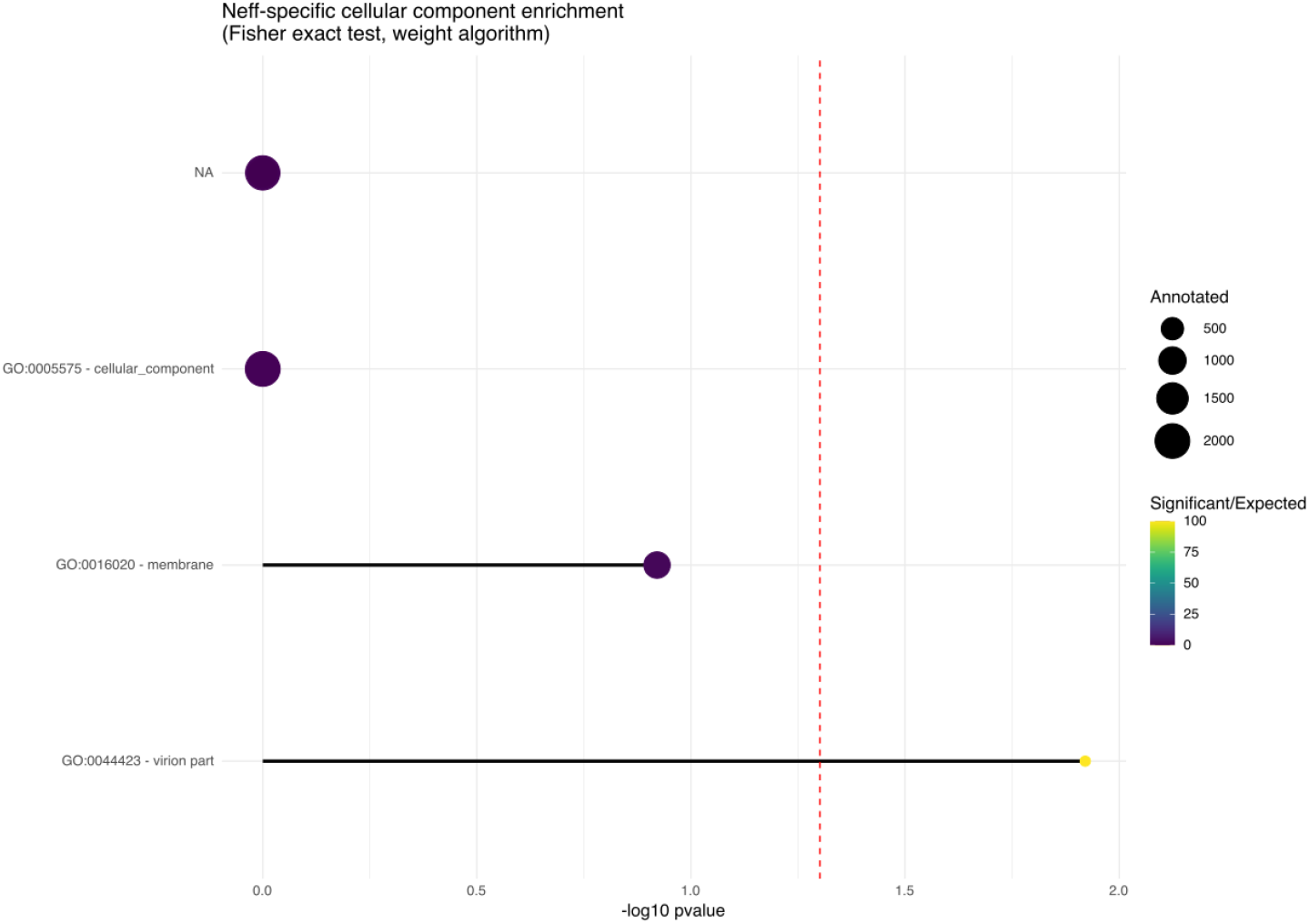
Most significant cellular component GO term enrichments in genes specific to Acanthamoeba castellanii strain Neff. Enrichment was determined using topGO, with nodeSize set to 5 when building the GOdata object. The size of the circles at the end of the bars represents the number of genes annotated under that GO term in the genome, and the colour scale of the circles represents the ratio of how many genes were found in the strain-specific set for that term compared to how many were expected.

**Figure S14.**
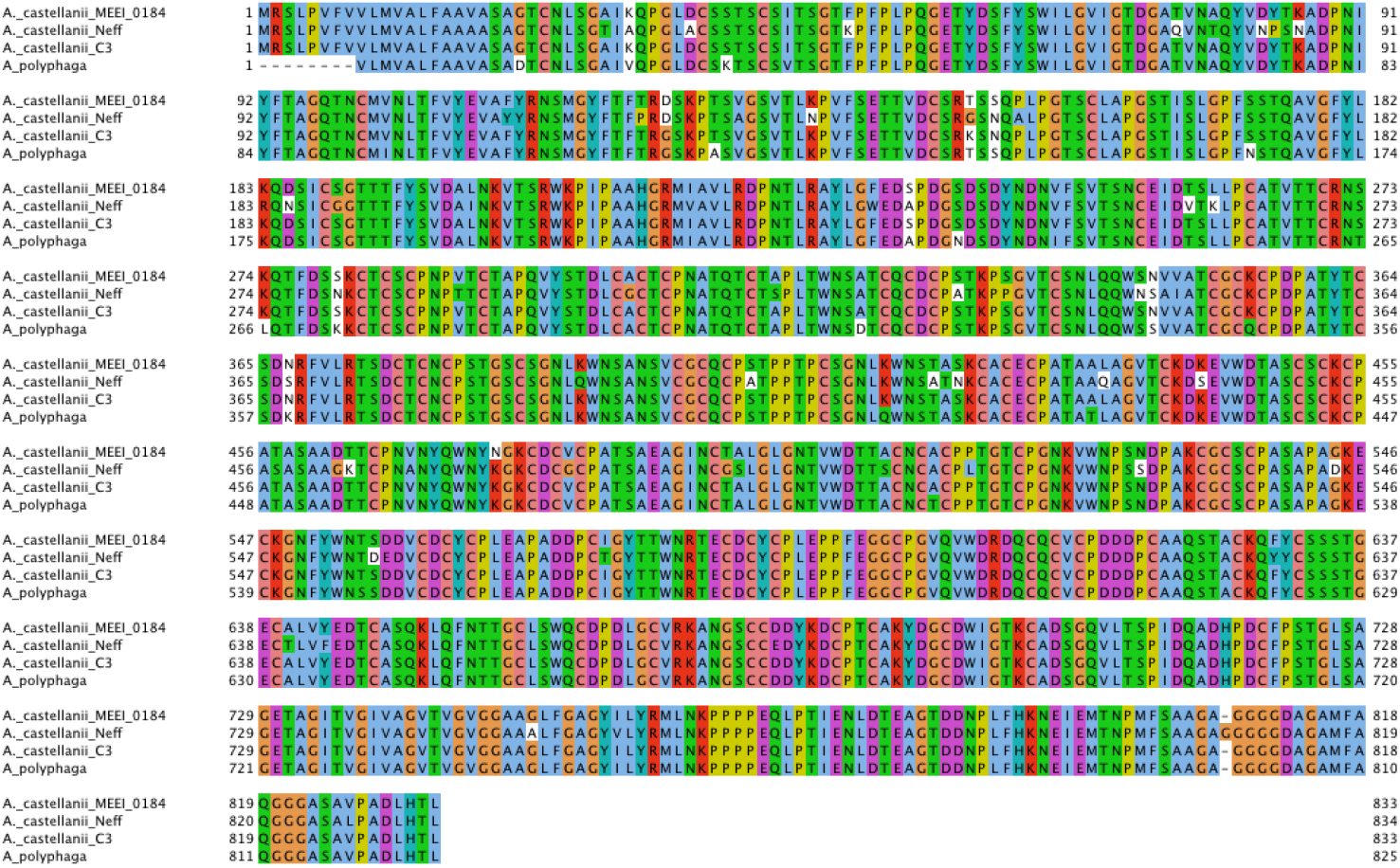
Multiple sequence alignment of mannose binding protein orthologs across three strains of Acanthamoeba castellanii and one strain of Acanthamoeba polyphaga. Sites are coloured according to the Clustalx colour scheme and residues differing from the consensus at any given site are not coloured. The alignment was generated with MAFFT-linsi, and was viewed and coloured in Jalview.

**Figure S15.**
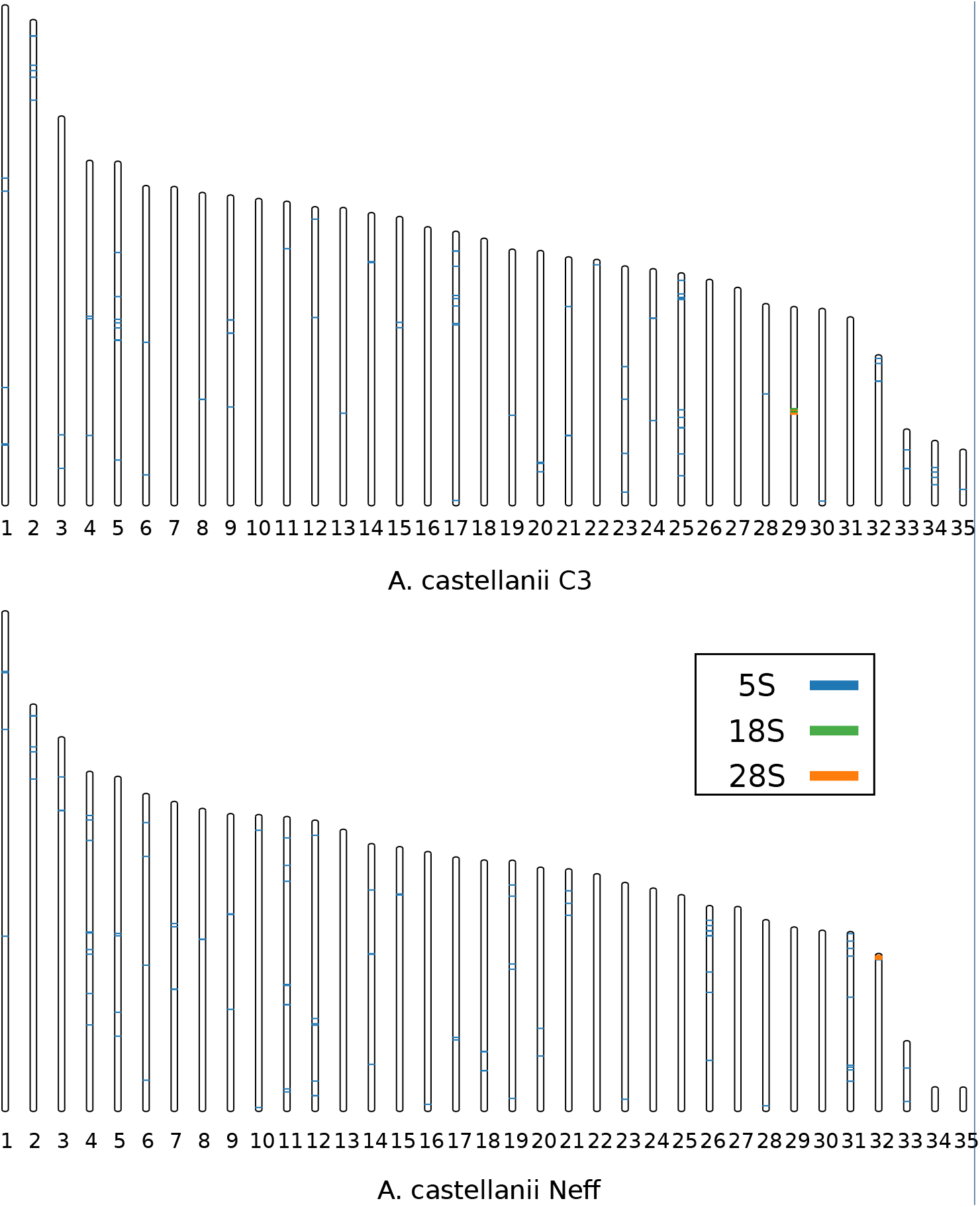
Predicted karyotype of A. castellanii strains C3 and Neff. For each strain, 35 scaffolds likely to be chromosomes based on the presence of inter-telomeric contact patterns on the contact maps are ordered by size. Colored bands indicate the position of rDNA along the chromosome sequence.

